# Cell type-specific responses to fungal infection in plants revealed by single-cell transcriptomics

**DOI:** 10.1101/2023.04.03.535386

**Authors:** Bozeng Tang, Li Feng, Pingtao Ding, Wenbo Ma

## Abstract

Plant infection by microbial pathogens is a dynamic process. Here, we investigated the heterogeneity of plant responses in the context of pathogen location. A single-cell atlas of *Arabidopsis thaliana* leaves challenged by the fungus *Colletotrichum* higginsianum revealed cell type-specific gene expression that highlights an enrichment of intracellular immune receptors in vasculature cells. Using trajectory inference, we assigned cells that directly interacted with the invasive hyphae. Further analysis of cells at these infection sites revealed transcriptional plasticity based on cell type. A reprogramming of abscisic acid signalling was specifically activated in guard cells. Consistently, a contact-dependent stomatal closure was observed, possibly representing a defense response that anticipates pathogen invasive growth. We defined cell type-specific deployments of genes activating indole glucosinolate biosynthesis at the infection sites, and determined their contribution to resistance. This research highlights the spatial dynamics of plant response during infection and reveals cell type-specific processes and gene functions.

## Introduction

Plant diseases are major threats to food security and global sustainability. To develop efficient disease resistance, it is crucial to have a comprehensive understanding of the molecular mechanisms involved in pathogen infection. However, pathogens infecting plant tissues are opportunistic and highly dynamic, and hence it is unclear how different plant tissues and cells respond to the pathogenic infection (Zhang et al., 2020). For instance, bacteria in plant phyllosphere were found to aggregate at preferred locations such as the base of glandular trichome, cell junctions, and veins (Lindow and Brandl, 2003). Bacterial entry into the leaf apoplast is often through stomatal openings formed by guard cells (McLachlan et al., 2014; Melotto et al., 2006). After entering the leaf tissue, the bacteria establish micro-colonies in close proximity to a subset of host cells (Badel et al., 2006), which may trigger distinct cellular responses compared to cells that are not directly targeted by the pathogen (Zhu et al., 2022).

The unequal interactions among plant cells with invading pathogens are more prominent during the infection of fungi and oomycetes, which undergo a sophisticated morphogenesis progress (Bozkurt and Kamoun, 2020; Fawke et al., 2015; Latijnhouwers et al., 2003; O’Connell et al., 2012; Wilson and Talbot, 2009). In this case, the heterogeneity in host responses is affected by both pathogen distribution and developmental changes. A typical infection process by a hemibiotrophic fungal pathogen begins with spore germination on the surface of plant tissue, followed by the development of appressorium. This specialised structure facilitates fungal penetration into host epidermal cells through plant cuticle (Wilson and Talbot, 2009). After penetration, the fungus develops invasive hyphae and establishes biotrophic growth inside the infected cell (Khang et al., 2010; Koeck et al., 2011; O’Connell *et al*., 2012). During the biotrophic infection stage, the fungal hypha is confined within the invaded cell, which remains alive and provides nutrients to the pathogen (Khang *et al*., 2010; McDowell, 2013; Ökmen and Doehlemann, 2014; Spanu et al., 2010). At a later infection stage, the pathogen switches to necrotrophic growth and feeds on the dead tissue. At the same time, invasive fungal hyphae move from the first infected cells into neighbouring cells, which support biotrophic fungal growth again (De Silva et al., 2017; Münch et al., 2008). Finally, the production and dissemination of fungal spores complete the infection cycle. The spatial context of pathogen localisation and the stage of infection and development can significantly shape the response of individual plant cells, highlighting the importance of taking spatial variability into account when investigating plant-pathogen interactions. Furthermore, plant immune responses may also exhibit cell type-specific features, adding to the complication of this highly dynamic process.

Plants have developed a highly sophisticated and, in most cases, robust immune system. The central components of plant immunity are immune receptors, which sense potential pathogens in the surrounding environment and activate the immune signalling (Dodds and Rathjen, 2010; Ngou et al., 2022). Cell surface-localized receptor-like kinases (RLKs) or receptor-like proteins (RLPs) recognise extracellular non-self-molecular signatures. They activate downstream molecular events, including ROS burst, Ca^2+^ influx, mitogen-activated protein kinase (MAPK) activation, and transcriptional reprogramming (Aerts et al., 2022; Ngou *et al*., 2022; Tsuda and Katagiri, 2010). Another class of immune receptors are intracellular nucleotide-binding domain leucine-rich repeat-containing receptors (NLRs), which sense pathogen effectors and activate effector-triggered immunity (ETI) (Cui et al., 2015; Ngou *et al*., 2022). NLRs in *Arabidopsis thaliana* (Arabidopsis hereafter) plants can be further classified based on their distinctive N-terminal domains as CC (coiled-coil)-NLR or CNL, TIR (Toll/Interleukin-1 receptor/Resistance protein)-NLR or TNL, and helper NLRs or RNL, which contain the CC_R_ domain (Gong et al., 2023; Kourelis et al., 2021). How the expression and function of RLKs/RLPs and NLRs are differentiated in different cell types during pathogen infection is largely unknown.

To investigate the transcriptional heterogeneity in plant response to fungal infection, we employed the droplet-based single-cell RNA-seq (scRNA-seq), which enables parallel transcriptome profiling of individual cells from a single reaction (Mazutis et al., 2013). RNA-seq has been extensively applied to understand plant-microbe interactions (Birkenbihl et al., 2017; Nobori et al., 2018; Tsuda and Somssich, 2015), through which many key molecular events were identified. However, bulk RNA-seq cannot capture differences between cell populations having specific interactions with the invading pathogen. Other methods, such as laser-microdissection (Chandran et al., 2010; Tang et al., 2006) and fluorescence-activated cell sorting-based enrichment (Rich-Griffin et al., 2020) can provide spatial information but are not high throughput. ScRNA-seq offers a new opportunity to understand the heterogeneity in gene expression changes among different cell populations during pathogen infection with a high resolution. In the past few years, a toolkit has been developed that enables comprehensive analyses of scRNA-seq data, further empowering the analysis of progressive processes (Saelens et al., 2019).

In this study, we determined cell type-specific responses of Arabidopsis to the infection of the hemibiotrophic fungal pathogen *Colletotrichum higginsianum*. We generated a transcriptome atlas constituted of 95,040 cells in the leaf tissue during biotrophic infection. Our dataset covered all major cell types, allowing us to reveal extensive heterogeneity in the expression of immune receptor genes in specific cell types. We combined quantitative live-cell imaging and trajectory inference to define the spatiotemporal dynamics of cellular response in the infected sites. We identified cell type-specific processes and gene functions in response to fungal infection. This work demonstrates how novel molecular events and immune-related genes can be identified by dissecting the spatiotemporal dynamics during infection using single-cell-based analyses. The transcriptome atlas provides a valuable resource for further investigations of gene expression in the specific cell population(s) and novel molecular mechanisms that underlie host-pathogen interaction. This knowledge is instrumental for developing resistant crops that precisely express defence-related genes at the infection sites to minimise growth penalty.

## Result

### Construction of an Arabidopsis leaf cell transcriptome atlas during *C. higginsianum* infection

We inoculated 14-day Arabidopsis seedlings with a conidia suspension of *C. higginsianum* and monitored the infection progression using live cell imaging. Extensive unevenness of fungal distribution in the inoculated tissue was observed, with only a portion of the cells in direct contact with the fungus at various developmental stages (Movie 1). Based on the microscopy analysis, two time points, 24 and 40 hours post inoculation (hpi), were selected for tissue collection as they represented early and late biotrophic stages, respectively (Fig. 1A). Protoplasts isolated from these tissues were subjected to single-cell partitioning and RNA library construction using the 10xGenomics workflow. The sequencing reads were aligned against the Arabidopsis genome (Table S1). After quality control and removal of batch effects, we generated a dataset that includes gene expression profiles of 95,040 cells (Fig. S1A).

**Figure 1.**
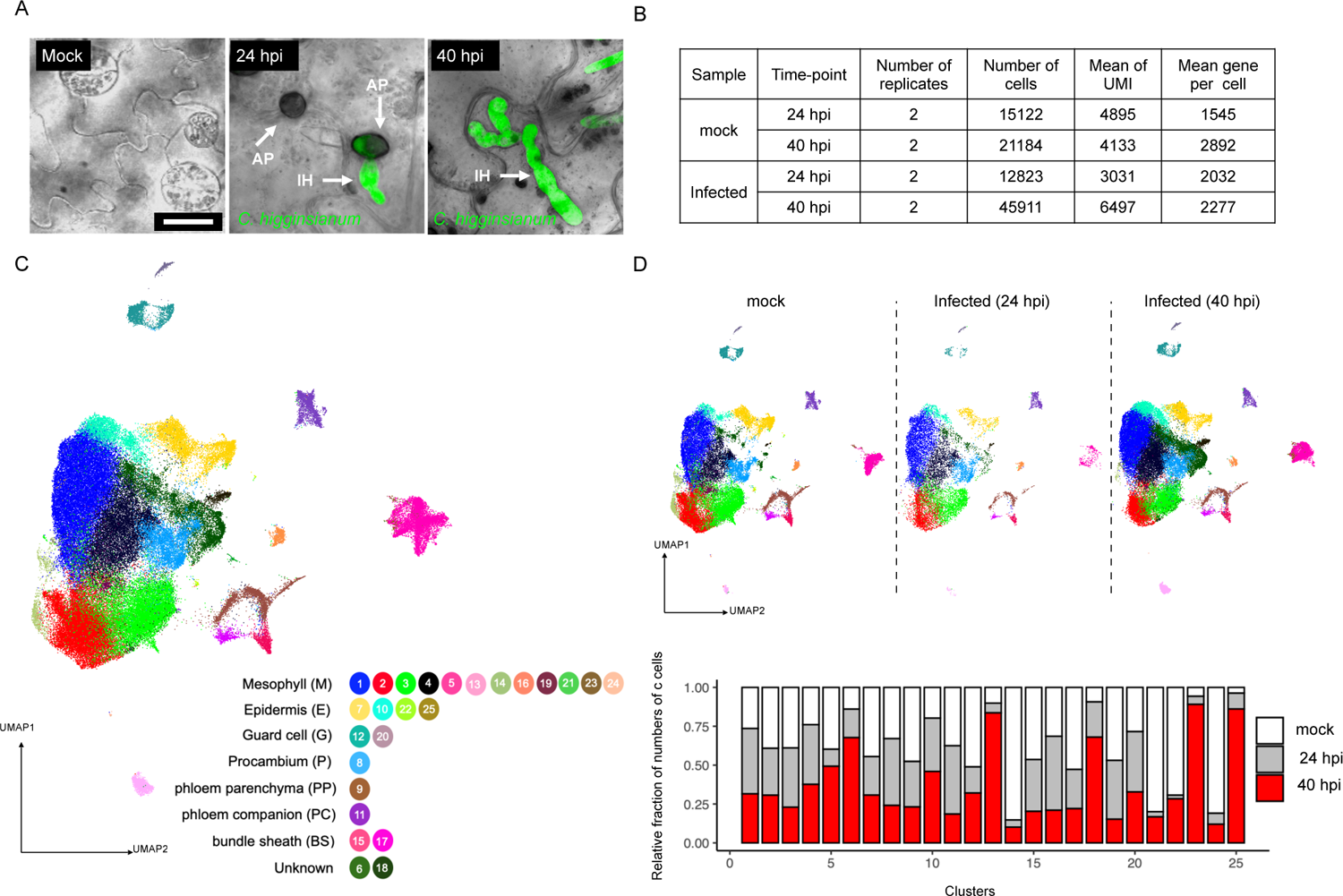
Construction of an Arabidopsis leaf cell transcriptome atlas during fungal infection. **(A)** Micrographs of Arabidopsis seedlings inoculated with conidia suspensions of *Collectotrichum higginsianum* showing uneven distribution of the fungus in leaf tissue. The fungus constitutively expressed green fluorescence protein (GFP) to facilitate visualization. Images were taken at 24 hours and 40 hours post inoculation (hpi), showing the progression of fungal invasion. Appressoria (AP) and invasive hyphae (IH) are labelled by arrowheads. Mock = water treatment. Scale bar = 10 µm. **(B)** A summary of the scRNA-seq data using leaf tissues at 24 and 40 hours after mock treatment or fungal inoculation. Unique Molecular Identity (UMI) = transcripts detected per cell. **(C)** A single cell atlas consisting of 95,040 cells. Each cell is represented by a dot on a plot visualized by uniform manifold approximation and projection (UMAP). Based on transcriptional signatures, the cells were divided into 25 clusters using graph-based unsupervised clustering. Each cluster was assigned to a cell type except for clusters 6 and 18. **(D)** Transcriptome changes in response to fungal infection captured by scRNA-seq. Upper panel: separate single cell atlas generated from mock treated and infected tissues (24 and 40 hpi). The 25 clusters described in (**C**) are presented in each atlas. Lower panel: proportions of cells from mock, 24 hpi or 40 hpi samples in each cluster. The proportions were normalized against the total number of cells in each sample.

On average, 2,583 genes and 5,491 Unique Molecular Identifier (UMI) were detected per cell, which covered 23,809 genes, representing > 90% of all protein-encoding genes in Arabidopsis (Fig. 1B). We then generated a single-cell transcriptome atlas using a graph-based unsupervised clustering analysis, which divided the cells into 25 clusters (Fig.1C), with cell numbers ranging from 20,500 (cluster 1) to 31 (cluster 25). Using cluster-enriched genes (Fig. S1B, Table S2) and reported marker genes (Chen et al., 2021; Kim et al., 2021; Lopez-Anido et al., 2021; Xu et al., 2022) (Fig. S1C), we were able to assign a specific cell-type identity to each cluster except for clusters 6 and 18 (Fig. 1C and S1D). Eleven clusters, including the top four clusters with the largest number of cells, were mesophyll cells, representing 70% of the total cells. They were followed by vasculature (six clusters, 13% of the total cells), the epidermis (four clusters, 8% of the total cells), and guard cells (two clusters, 3% of the total cells).

We next analysed the proportions of cells in each cluster from the different treatment samples. In this analysis, samples collected 24 and 40 hours after mock (water) treatment were merged because they exhibited highly similar gene expression patterns based on a Pearson correlation analysis (R^2^ > 0.91). We found that the fractions of cells from different treatments can be drastically different in some clusters. In particular, clusters 6, 13, 18, 23 and 25 mainly constitute cells from the 40 hpi sample (Fig. 1D), while clusters 14, 21 and 24 mainly constitute cells from the mock-treated samples. Interestingly, a clear cell type identity could not be assigned to clusters 6 and 18, possibly due to a global transcriptome reprogramming in response to fungal infection. Similarly, atlases generated by cells from each treatment also showed significant changes in specific cell clusters at 40 hpi (Fig. 1D).

To eliminate the possibility that the analysis might be skewed by the protoplasting process during sample preparation, we determined gene expression changes caused by protoplast isolation using bulk RNA-seq. We found that protoplasting-induced transcripts, as well as mitochondrial and chloroplast genes, which may also reflect potential cell damage, were generally lower than 10% in each cluster and this proportion was relatively constant crossing the clusters (Fig. S1E). This result demonstrates that the specific transcriptome signatures and the clustering of cell populations were predominantly determined by cell-type determinants and cellular responses to fungal infection. Taken together, our analysis generated a single-cell atlas covering all major cell types in Arabidopsis leaf tissue and captured dynamic gene expression changes during fungal infection in different cell populations.

### Cell type-specific expression of genes encoding immune receptors

We examined differential gene expression in specific cell types, some of which may have been masked in bulk RNA-seq analyses. A comparison to a previous bulk RNA-seq study using the same Arabidopsis-*C. higginsianum* plant-pathogen interaction system (O’Connell *et al*., 2012), strongly correlated with our scRNA-seq data on genes differentially expressed after fungal infection. However, many genes considered to have an unchanged expression in bulk RNA-seq showed differential expression in a cell type-specific manner (Fig. 2A).

**Figure 2.**
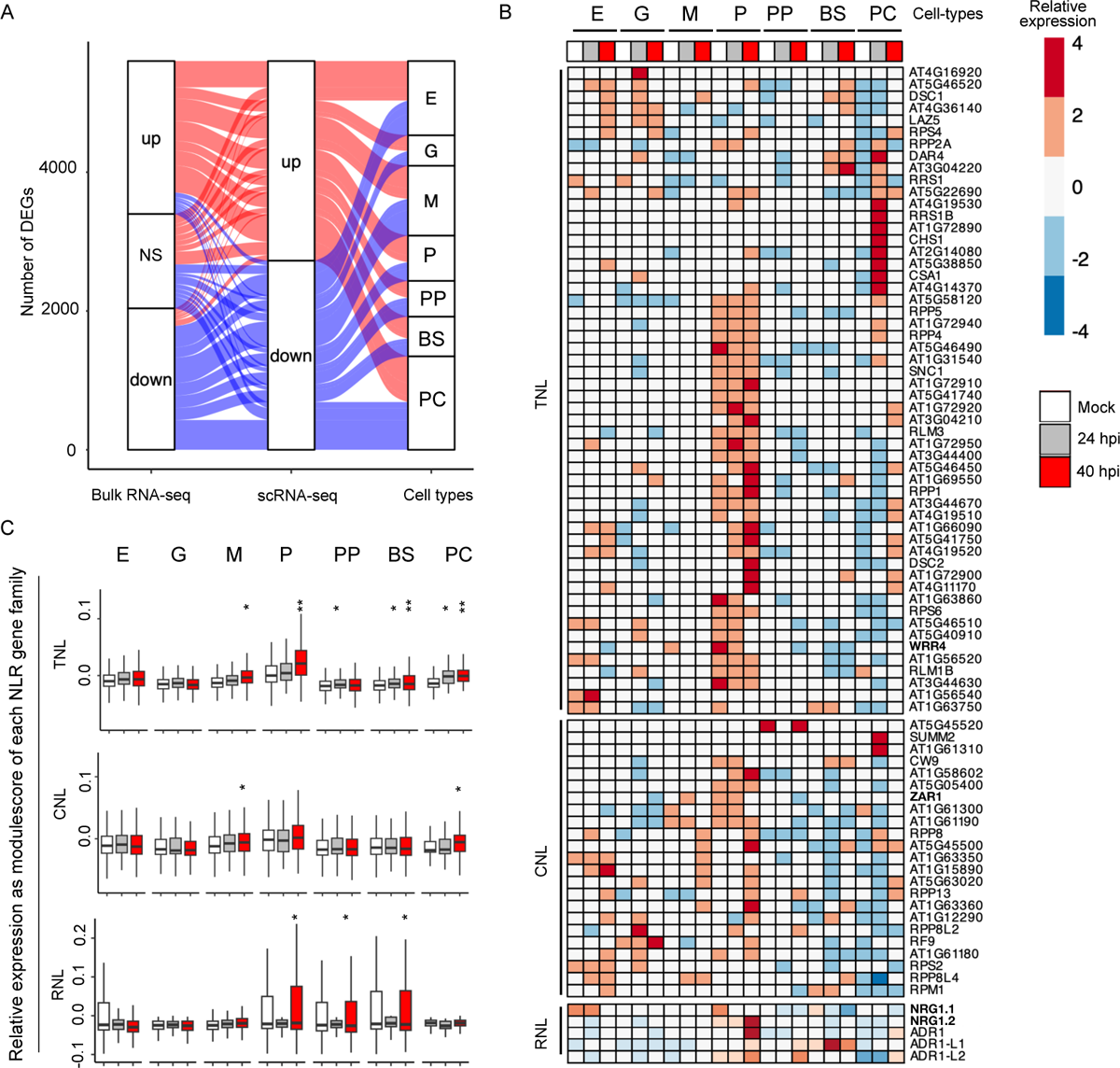
Cell type-specific expression of immune receptors in Arabidopsis. **(A)** ScRNA-seq captured cell type-specific changes in gene expression after fungal infection. Differentially expressed genes (DEGs) identified from scRNA-seq was compared with DEGs detected from a published dataset (ENA projects PRJNA148307, PRJNA151285, and PRJNA438701) using bulk RNA-seq. The group “NS” includes genes that were not significantly changed in their expression in bulked RNA-seq but showed differential expression in a cell type-specific manner based on scRNA-seq. DEGs calling from the scRNA-seq data was done by FindMarker with default settings in each annotated cell type. DEGs calling from the bulk RNA-seq used cut-offs of log_2_|FC| >1, p-adj <0.05. **(B)** Cell type-specific expression and induction by fungal infection of NLRs. Heatmaps show relative expression of known or predicted NLR-encoding genes in each annotated cell type and with or without fungal infection. NLRs were predicted from the Arabidopsis genome using NLRtracker, which include 70 TNLs, 23 CNLs, and 5 RNLs. Gene expression levels are represented by aggregated UMI counts in all the cells belonging to the same cell type and in the same treatment. **(C)** Specific induction of TNLs in vasculature cells after fungal infection. Boxplots show normalized expression of all genes belonging to the same NLR family from different cell types. The relative expression of an NLR was normalized to 50 randomly selected genes in the same cell. The normalized expression levels of genes belonging to the same NLR family in the same cell type were then aggerated and compared between mock treated and fungal infected samples. T-test was used to determine the significant differences. * = *p* <0.05, ** = *p* <0.01.

Given the essential role of immune receptors in plant defence, we investigated their expression patterns in different cell types. We predicted 170 genes from the Arabidopsis genome that putatively encode the intracellular immune receptors NLRs using NLRtracker (Kourelis *et al*., 2021). Ninety-eight of these predicted NLR genes were detectable from the transcriptome of at least 500 cells (Table S3). These genes were further analysed by determining their relative expression levels in each cell type in mock and fungal infection samples. Interestingly, this analysis revealed that a significant portion of TNL genes, including those encoding TNLs with known functions such as *RPP1*, *RPP5*, *SNC1*, and *RPS6*, were predominantly expressed in the procambium cells with either mock treatment or fungal infection (Fig. 2B). Procambium is a cell type from which primary phloem and xylem cells derive (De Rybel et al., 2016; Jouannet et al., 2015). Many of these procambium-expressed *NLR* genes were further induced after fungal infection (Fig. 2B). In addition, another group of *TNL* genes are highly, and almost exclusively induced in phloem companion cells at 24 hours but not at 40 hours post fungal inoculation (Fig. 2B). These include genes encoding TNLs with known functions such as *RRS1B*, *CSA1* and *CHS1*. This pattern of transient induction may indicate a suppression from the pathogen of host immunity. A small number of *TNL*s were induced by fungal infection in multiple cell types. For example, *RPS4* showed increased expression in epidermal, guard, procambium, and companion cells.

In contrast to TNLs, a general trend of vasculature-specific expression was not observed in CNLs or RNLs except for in individual cases (Fig. 2B). For example, a cluster of CNL genes including *CW9* and *ZAR1* showed relatively higher expression in vasculature cells; *RF9* was predominantly expressed, and further induced by the fungal infection, in guard cells; and the transcripts of *SUMM2* was only detected in companion cells at 24 hpi. Similar to CNLs, RNLs did not exhibit a specific expression pattern in a particular cell type. However, genes in the same family showed differences in their expression in different cell types. For the three *ADR1* homologs, *ADR1* and *ADR1-L2* were specifically expressed in procambium and phloem cells. Their expression was also highly induced after fungal infection, especially at 40 hpi (Fig. 2B). In contrast, *ADR1-L1* had a specifically high expression in bundle sheath cells, which was also induced by fungal infection at both 24 and 40 hpi. For the *NRG1* homologs, *NRG1.1* was expressed in epidermal and procambium cells, whereas *NRG1.2* was mainly expressed in procambium cells. Interestingly, the expression of *NRG1.1* was decreased after fungal infection, especially at 40 hpi; however, *NRG1.2* was strongly induced after fungal infection. RNLs play an essential role in immune signalling. The strong pattern of cell type-specific expression indicates potential functional variation in these RNL homologs and suggest a fine-tuning in their deployment. The contrast expression changes in the NRG1 homologs may represent a defence-counter-defence arms race during fungal infection. For example, the pathogen may manipulate the expression of *NRG1.1* to suppress plant defense.

To detect general cell type-specific patterns in the expression changes of NLRs after fungal infection, we compared the aggregated expression of the three NLR families at 24 and 40 hpi to mock treatment in each cell type (Fig. 2C). A strong overall induction was only observed in TNL genes in vasculature-related cell types including procambium, bundle sheath, and companion cells. This is consistent with the vasculature-specific expression of many TNL genes, representing a previously unknown phenomenon in plant immunity. In addition, we also observed a strong correlation of the expression patterns between NLRs and RLKs in almost all the cell types after fungal infection (Fig. S2). This observation reflects a potential coordination among different classes of immune receptors.

### Spatial heterogeneity of gene expression in correlation to fungal distribution

In the transcriptome atlas, clusters 6 and 18 were special because: 1) they constitute cells predominantly from infected tissues (Fig. 1D, S3A); and 2) unlike the other cell clusters, cells in these two clusters could not be categorised into a specific cell type. Therefore, cells belonging to these two clusters have unique gene expression signatures, likely due to significant transcriptome reprogramming in response to fungal infection. Indeed, we found that these cells were enriched with transcripts from genes that were previously found to be induced by *C. higginsianum* infection using bulked RNA-seq (Poncini et al., 2017; Zhou et al., 2020) (Fig. S3B). Using all the cells belonging to these two clusters (6,603 in total) as one input, we conducted a sub-clustering analysis, which defined 25 sub-clusters and successfully assigned them to the epidermis, mesophyll, vasculature, or guard cells, respectively (Fig. S3C, Table S4). These cells were combined with other cells that have already been classified to each cell type to construct trajectory curves to dissect the progressive changes in the transcriptome of plant cells during infection in a cell type-specific manner.

Fungal infection is a spatiotemporal dynamic process with only a portion of plant cells in direct contact with the pathogen. A trajectory analysis aims to capture transcriptional signatures at the single-cell level that exhibit gradual changes. In our study, the trajectory may reflect variable interactions of plant cells with the fungal pathogen as a continuous process. Datasets from four cell types in tissues from both mock treatment and fungal infection were individually analysed using monocle (Cao et al., 2019), through which a pseudotime value was assigned to each cell to indicate their respective status. A trajectory curve was then generated to depict gradual changes in these cells during infection. By comparing the trajectory curves of all four cell types between mock treatment and fungal infection, we observed a shift towards pseudotime of “1” was revealed in the cell populations after fungal infection with cells from the 24 hpi sample enriched in the middle of the curve and cells having pseudotime values between 0.5 and 1 predominantly from the 40 hpi sample (Fig. 3A). This shift of cell populations with the progression of the fungal infection could also be demonstrated by comparing to trajectory curves generated by cells from 24 hpi, and 40 hpi samples with the mock treatment (Fig. 3B).

**Figure 3.**
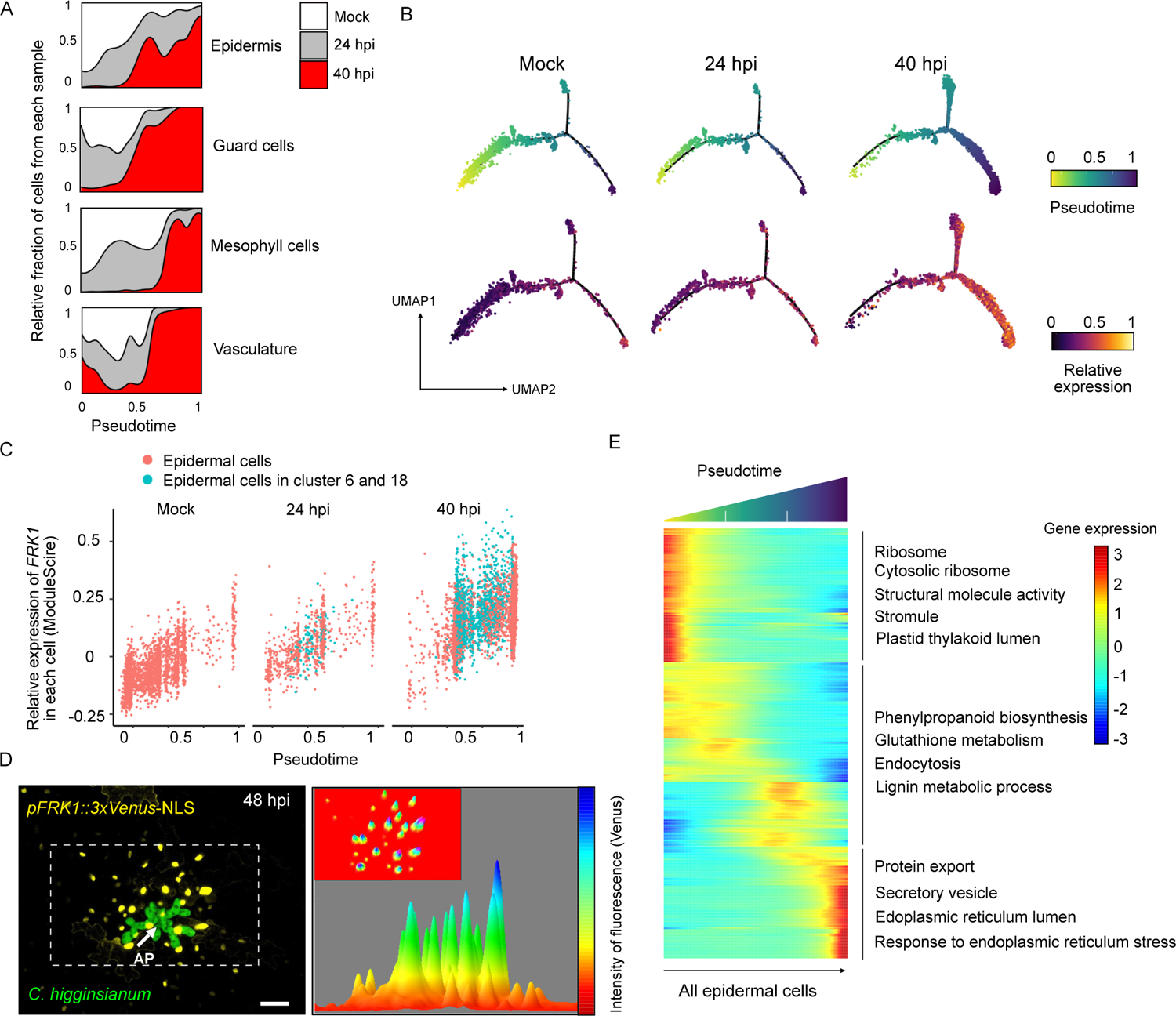
Spatiotemporal heterogeneity of cellular response to fungal distribution. **(A)** A shift towards pseudotime of “1” in the cell populations after fungal infection. The density plots show fractions of cells with different pseudotime values from mock, 24 hpi or 40 hpi samples in each of the four major cell types. Each cell was assigned a pseudotime value based on transcriptional signatures that exhibit gradual changes. **(B)** The cell population shift in response to fungal infection corelates with increased expression of infection-site-induced genes in these cells. Upper panel: trajectory curves of epidermal cells from mock, 24 hpi and 40 hpi samples with each dot representing a single cell. The colour of a dot represents its pseudotime value. Lower panel: aggregated expression level of a set of 41 genes that were identified to be induced at the infection site of powdery mildew (Chandran *et al*., 2010) mapped to the trajectory curves. The colour intensity of a dot indicates the relative value of module-score in the cell. **(C)** *FRK1* expression showed gradual increase in epidermal cells with pseudotime values from 0 towards 1. Each dot represents a cell. Blue dots represent cells from the cluster 6 and 18 in Figure 1D, which mainly consist of cells from the 40 hpi sample. These cells also had relatively higher levels of *FRK1* expression. **(D)** *FRK1* was highly induced in cells that had direct contact with invading fungus. Arabidopsis seedlings expressing 3xVenus-NLS under the *FRK1* promoter were inoculated with *C. higginsianum* and imaged at 48 hpi. A representative micrograph shows the fluorescence signal of Venus (yellow), which indicates the promoter activity of *FRK1*, and green fluorescence expressed in the fungal invasive hyphae. The arrowhead indicates the penetration site by fungal appressoria (AP). The boxed area was further analyzed by quantifying the Venus signal intensity, which is shown in the plot from a horizontal angle. The mini micrograph shows the same distribution of Venus signals but from a vertical angle. Scale bar = 15 µm.**(E)** Different cellular processes enriched in cell populations with different pseudotime values. The Heatmap shows relative expression levels of pseudotime-dependent genes in the epidermal trajectory. The cells were grouped into three populations based on a hierarchical cluster analysis. Representative enriched KEGG metabolic pathways and GO terms identified in each cell population are highlighted.

To interpret the biological meaning of pseudotime, we analysed the expression of a set of 41 genes (Table S5), which were previously identified to be induced at the infection sites after powdery mildew infection using a laser-microdissection method (Chandran *et al*., 2010). We observed a strong correlation between the induced expression of these genes and pseudotime values, indicating that an increase in gene expression led to a corresponding increase in pseudotime value within each cell (Fig. 3B, S3D). This result indicates that the pseudotime value of each cell reflects its relative position to the infection sites. To further support this correlation, we mapped the transcript level of *FRK1*, one of the 41 genes analysed above, in each epidermal cell in 24 hpi, and 40 hpi cell populations with the mock treatment. A clear trend of gradual increase in *FRK1* expression was observed in cells when their pseudotime values increased from 0 towards 1 (Fig. 3C). The cells with relatively higher expression of *FRK1* were particularly enriched in the 40 hpi sample. Importantly, cells belonging to clusters 6 and 18 also showed significant enrichment of pseudotime values between 0.5 and 1 in the 40 hpi sample and exhibited high *FRK1* expression (Fig. 3C). These results confirmed that the cluster 6 and 18 cells were tightly associated with fungal infection. It is also noteworthy that including all cells belonging to these two clusters was essential for generating the trajectory curve.

Next, we experimentally confirmed the correlation between the pseudotime values and the relative position of the cells with the invading fungus. Leaves of Arabidopsis transgenic plants expressing *pFRK1::3xVenus-*NLS (Poncini *et al*., 2017) were inoculated with conidial suspensions of *C. higginsianum* and subjected to live-cell imaging. The expression of *FRK1* was monitored by quantifying the intensity of yellow fluorescence at 48 hpi. *C. higginsianum* was labelled by green fluorescence, which was used to indicate the location of the fungus. The yellow fluorescence signals were specifically accumulated in the cells with direct contact or in close proximity to the fungal hyphae but were absent in more distal cells (Fig. 3D). Importantly, the strongest yellow fluorescence was observed from cells directly colonised by the fungus and the fluorescence levels gradually decreased in the neighbouring cells (Fig. 3D). This experiment confirmed that the dynamic cellular responses represented in the trajectory curve are indicative of a spatial heterogeneity determined by the uneven distribution of the fungal pathogen in the infected tissue.

Analysis of the transcriptional signatures in cells in the epidermal trajectory revealed 767 genes that showed pseudotime-dependent changes (Table S6, Fig. 3E). Among them, 296 genes were highly expressed in the cells at the beginning of the curve, none of which has known functions related with defence response. These cells were, therefore, likely distal to the invasive hyphae and not significantly affected by the infection. 329 genes were induced in cells in the middle of the curve, which may be surrounding but not located at the infection sites. Some of these genes encode positive regulators of plant defence, such as *KTI1*, *GSTU10*, and *WRKY75*, and their induction suggested activation of the immune response. Another 142 genes were specifically induced in cells at the end of the curve, which likely had direct interactions with the invasive hyphae. These also include well-studied immune-related genes such as *FRK1* and *PEP2*. Enrichment analysis demonstrates that distinctive cellular processes were activated in the different cell populations along the trajectory curve. For example, genes in secretion system-related pathways were specifically induced in cells at the infection sites (Fig. 3E), consistent with their direct involvement in anti-microbial activities. These analyses revealed major cellular processes in response to fungal infection in different plant cell populations based on their relative location with the pathogen.

### A guard cell-specific response leads to stomatal closure at the infection sites

Similar to the epidermal cells, the trajectory of guard cells also demonstrated a transcriptome switch in cells with pseudotime values towards 1 after fungal infection, which is in correlation to the induced expression of infection site-related genes (Fig. 4A). By investigating pseudotime-dependent genes, we discovered metabolic pathways associated with propionate metabolism, nitrogen metabolism, and sulfur metabolism, that were specifically enriched in guard cells at the infection sites (Fig. 4B). Sulfur metabolism is well known to be responsive to abscisic acid (ABA), which modulates stomatal closure (Batool et al., 2018). This led us to investigate the expression changes of ABA-related genes in the guard cells. A total of 104 ABA-related genes in *Arabidopsis* genome were pulled out for expression analysis, including 51 positive regulators and 53 negative regulators (Table S5). Strikingly, we observed a strong corelation between gene expression pattern of positive regulators and the pseudotime values in the guard cells. This is mainly primed by the genes as, such as *PED1*, *CAR8*, and *CPK11*, which showed a continuous increase in their expression in guard cells with pseudotime values towards 1 (Fig. 4C), indicating a gradual induction at the infection sites. Interestingly, an opposite pattern, i.e. gradual decrease, was observed in the expression of genes encoding negative regulators of ABA signalling, such as *ABR1*, *ABI1*, *FER1*, and *GCR1* (Fig. 4D). These expression patterns were specific to guard cells, as their expression in the other three cell types remained unchanged (Fig. S4). We, therefore, hypothesised that ABA signalling was activated in response to fungal infection in a spatially dynamic and cell type-specific manner.

**Figure 4.**
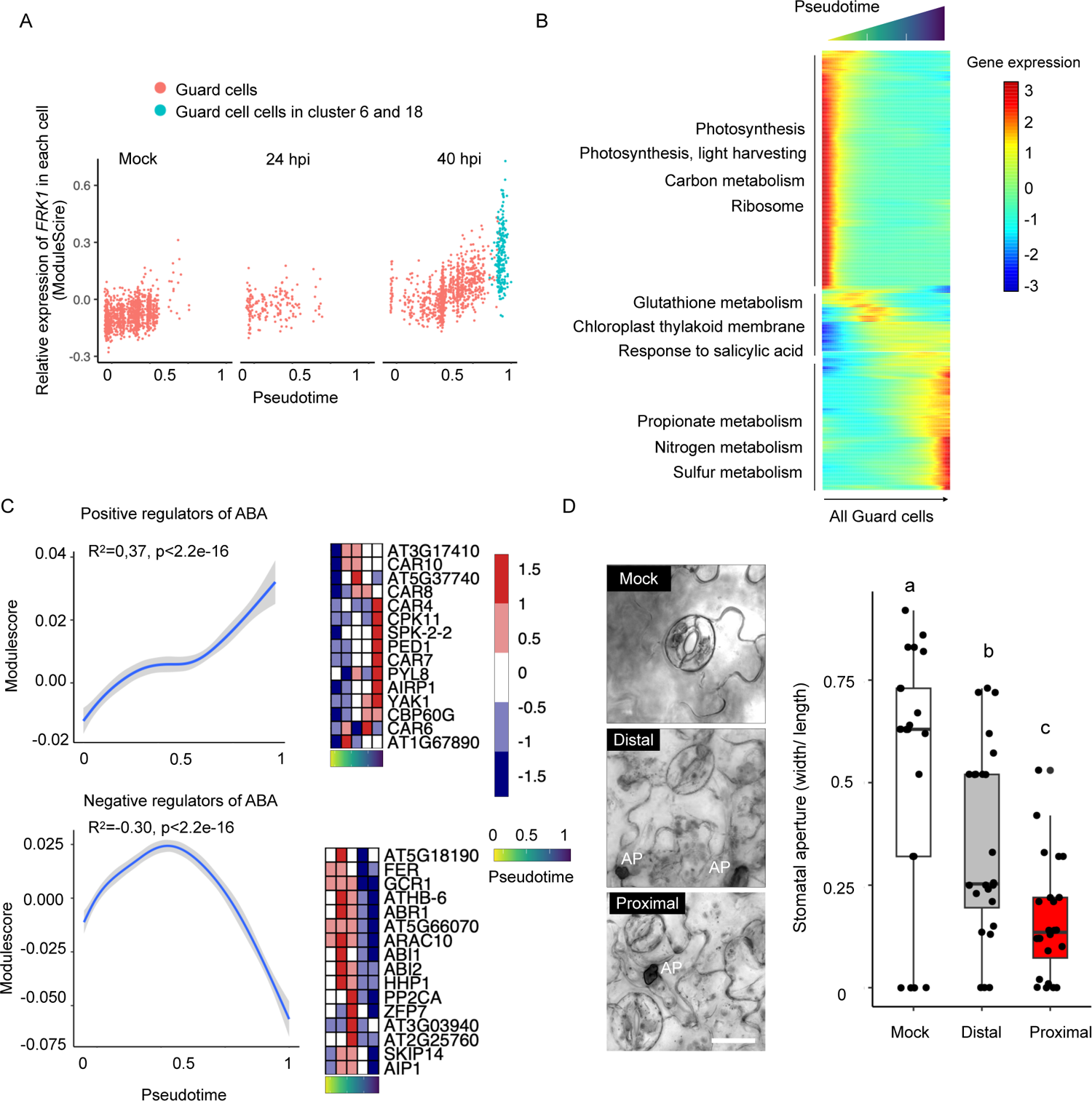
A Guard-cell-specific response leads to stomatal closure at the fungal infection site. **(A)** Gradual increase of *FRK1* expression in guard cells with pseudotime values from 0 to 1. Each dot represents a cell. Blue dots represent cells from the clusters 6 and 18 in Figure 1D. **(B)** Different cellular processes enriched in different guard cell populations after fungal infection. The Heatmap shows relative expression levels of pseudotime-dependent genes along the trajectory. Representative enriched GO terms are highlighted in each of the three cell populations. **(C)** ABA signalling activated in guard cells with pseudotime values toward “1”. The line plots show linear regression of the relative expression levels of genes encoding positive (upper panel) or negative (lower panel) regulators of ABA signalling in guard cells with their pseudotime values. The heatmaps show the expression profiles of genes that exhibited the most significant changes. **(D)** Stomata closure at the fungal infection sites. Representative micrographs from live-cell imaging show the status of stomatal aperture from the leaves treated with water (uninfected) or inoculated by *C. higginsianum* at 40 hpi. The “distal” guard cells were defined as being separated from the fungus by at least one epidermal cell but up to five cells. The “proximal” guard cells were defined as being located immediately adjacent to the appressoria. Scale bar =10 µm. Stomatal aperture was measurement using the micro-images of the guard cells. One-way ANOWA was used to determine the significant differences between the samples, which were labelled with different letters. The imaging acquisition was performed for at least three times for each experiment.

ABA is a major regulator of stomatal closure, which is important in preventing bacterial entry into the leaf apoplast (Hu et al., 2022; Lovelace and Ma, 2022; Roussin-Léveillée et al., 2022). We then examined whether the specific activation of ABA signalling led to changes in the stomatal aperture during fungal infection. For this purpose, confocal imaging was conducted on Arabidopsis leaf tissues inoculated with *C. higginsianum* to capture images of guard cells at 40 hpi. The width/length ratio was measured for each stomata using a machine learning-based imaging acquisition workflow (Sai et al., 2022) to quantify stomatal aperture. We measured the stomatal aperture in guard cells at the infection sites, in the more distal regions but still the neighbouring tissue of the invasive hyphae, and in mock-treated tissues. A comparison between these different guard cell populations revealed a decrease in the stomatal opening when the guard cells were in the proximity of the invading pathogen and complete closure when the cells were in direct contact with the fungus (Fig. 4E). These results show a guard cell-specific potentiation of ABA signalling, mediated by the coordinated expression changes of positive and negative regulators, that results in the closure of stomata as a potential defence response in anticipation of pathogen penetration.

### Activation of indole glucosinolate biosynthesis at infection sites employs cell type-specific genes

When comparing the cells at the infection sites to the whole atlas, we found that only five genes were induced in all four cell types. In contrast, 184, 193, 6, and 23 genes were specifically induced in epidermal, stomata, mesophyll, and vasculature cells, respectively (Fig. 5A, Table S7). For instance, NHL25 and CAD1 as components depend on the SA pathway were specifically induced in epidermal and guard cells, respectively (Fig. 5B). These genes in the above four cell types include the vast majority of genes identified from the previous laser-microdissection-based studies, but many additional infection site-related genes were revealed from our single cell-based analysis, which showed cell type-specific induction (Fig. S5A). Despite the strong pattern of cell type-specific transcription changes, enrichment analysis showed that indole glucosinolate (IG)-related pathways were induced in all cell types at the infection sites (Fig. S5B). IGs are defence-related metabolites that have also been found to be induced after pathogen infection (Frerigmann and Gigolashvili, 2014; Widemann et al., 2021; Xu et al., 2016). Interestingly, not all IG biosynthesis-related genes were induced in all cell types (Fig. S5C). This prompted us to examine the expression of genes with known functions in the IG biosynthetic pathway in the four cell types. We found that, except for *MYB51*, *ASA1* and *TSB1*, and *SOT16*, which had a general induction in all cell types, all other IG-related genes exhibited cell-type specificity in their expression (Fig. 5C). For instance, GGP1 and SUR1 both catalyse the conversion of *S*-alkyl-thiohydropximate to thiohydroximate; however, *GGP1* was induced only in epidermal cells while *SUR1* was induced in the other three cell types. Similarly, *CYP82F1*, *CYP81F2* and *CYP81F3* all catalyse the conversion from indole-3-ylmethyl-glucosinolates (13M) to 4-hydroxy-indole-3-ylmethyl glucosinolates (4HO-13M) but exhibited different cell type-specific expression: *CYP82F1* was induced in epidermal cells, *CYP82F3* was induced in mesophyll cells, and *CYP82F2* was induced in the mesophyll, vasculature, and guard cells (Fig. 5B).

**Figure 5.**
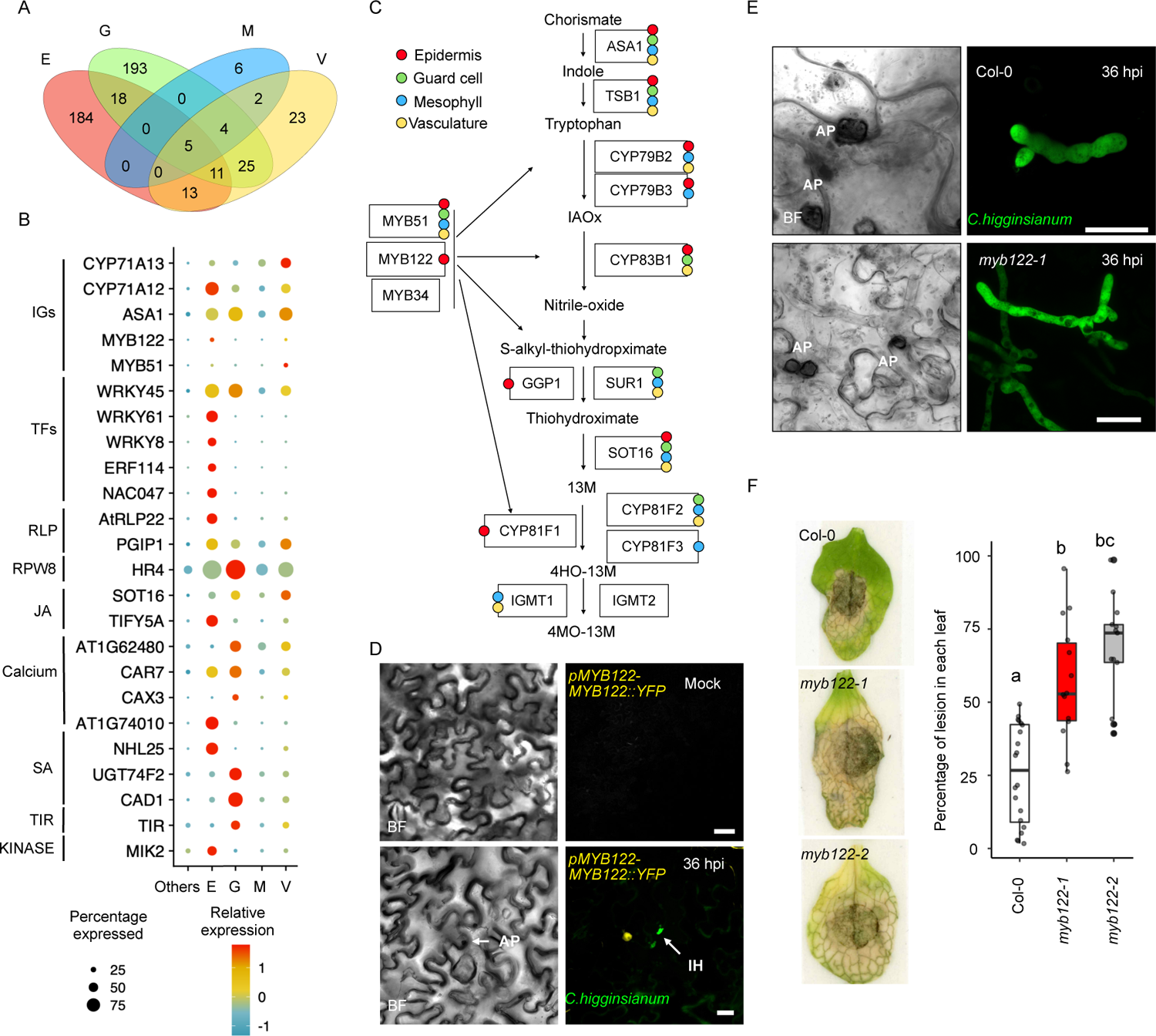
*MYB122* was specifically induced in epidermal cells at the infection sites and contributed to fungal resistance. **(A)** Cell-type-specific induction of genes in cells that had direct contact with the invading fungus. The Venn-diagram shows the number of induced genes identified from each of the four major cell types at the fungal infection sites. The cut-off was set as Log_2_|FC| >1.5, p-adj <0.05.**(B)** Genes specially induced in infection site cells. Expression profiles of genes of interest show specific induction at the infection sites in four major cell types. The size of the dots indicates the proportion of cells in which the corresponding gene was induced, and the colours indicate relative expression levels in the cell. **(C)** Cell-type-specific induction of genes involved in indole glucosinolates (IG) biosynthesis. Arabidopsis genes with proposed functions in the IG biosynthetic pathway (Frerigmann and Gigolashvili, 2014) were analyzed for cell-type-specific activation at the infection sites. **(D)** Specific induction of *MYB122* in the epidermal cells that had direct contact with the invading fungus. *Nicotiana benthamiana* leaves transiently expressing *pMYB122*::*MYB122-YFP*. At 24-hour post-Agroinfiltration, the leaves were inoculated with conidia suspension of *C. higginsianum* or water (mock). Images were taken at 36 hpi. Yellow fluorescence signals in the nucleus represent MYB122 expression and the green fluorescence signals represent fungal invasive hyphae. Arrowheads point to fungal appressoria (AP) and invasive hyphae (IH). Scale bar = 30 µm.**(E)** Accelerated progression of fungal infection in an *MYB122* mutant of Arabidopsis. Micrographs show *C. higginsianum* hyphae development in the leaves of wild-type (Col-0) or a *MYB122* mutant (*myb122-1*) plants at 36 hpi. Green fluorescence signals represent fungal invasive hyphae. Fungal appressoria (AP) are labelled. Scale bar = 10 µm.**(F)** Hypersusceptibility of *MYB122* mutants to *C. higginsianum*. Two T-DNA insertion mutants of *MYB122* were analysed for susceptibility compared to wild-type Arabidopsis (Col-0). Leaves of four weeks old Arabidopsis plants were inoculated by conidial suspension of *C. higginsianum*. Disease symptoms were monitored at 5 days post-inoculation. The lesion size of each leaf was also analysed as the ratio of discoloured area vs total leaf area. Each experiment was repeated at least three times. One-way ANOVA was used to determine the significant differences between the samples, which were labelled with different letters (*p* < 0.05).

We were particularly interested in the transcription factor *MYB122*, specifically activated in epidermal cells at the infection sites. MYB122 is one of the three transcription factors proposed to regulate IG biosynthesis by activating the expression of several key enzymes in the pathway. For genes encoding the other two transcription factors, *MYB51* was induced in all four cell types, while *MYB34* was not expressed in any cell types at the infection site. Unlike MYB51, which is known to be defence responsive (Zhou *et al*., 2020), MYB122 was not previously known to play a role in plant immunity. To confirm the specific *MYB122* induction at the epidermal infection sites, we transiently expressed *MYB122-YFP* driven by the native promoter of *MYB122* in *Nicotiana benthamiana* and then inoculated the *Agrobacterium*-infiltrated leaves with *C. higginsianum*. At 24 hours post fungal inoculation, we observed yellow fluorescence signals representing MYB122 expression only in the *C. higginsianum*-inoculated tissues and, more specifically, in cells in close proximity to the fungal appressoria and invasive hyphae (Fig. 5C). We next determined the functional relevance of *MYB122* induction using two independent Arabidopsis mutants with T-DNA insertions in *MYB122* gene-coding region. We then performed live-cell imaging to monitor fungal invasive growth in the epidermal cells of Arabidopsis leaves inoculated with conidial suspensions of *C. higginsianum* at 36 hpi (Fig. S5D). We observed accelerated biotrophic growth of the fungus in the epidermal cells in the *myb122* mutants wild-type leaves compared to that in wild-type (Fig. 5D). Consistent with this observation, the *myb122* mutants exhibited enhanced susceptibility to *C. higginsianum* as shown with larger lesion sizes and more severe disease symptoms at 5 days post inoculation (dpi) (Fig. 5E). These results demonstrate a significant contribution of MYB122 in plant defence, presumably through elevating the production of IGs specifically in epidermal cells at the infection sites, which restricts fungal biotrophic growth. These findings emphasise the recruitment of different sets of genes involved in the induced IG biosynthesis in different cell types.

## Discussion

Single cell-based omics analyses have been proven to be powerful tools in constructing the progression of a biological process and understanding the heterogeneity within different cell populations. In recent years, this technology has been applied to plant science, especially in many developmental processes (Kim *et al*., 2021; Liu et al., 2020; Lopez-Anido *et al*., 2021; Shahan et al., 2022), and more recently, plant-microbe interactions (Cervantes-Pérez et al., 2022; Illouz-Eliaz et al., 2023; Ye et al., 2022; Zhu *et al*., 2022). Here, we generated a single-cell transcriptome atlas with ∼95,000 cells during the infection of a fungal pathogen that colonises the leaf tissue of the model plant Arabidopsis. This atlas captured all major cell types in leaves, allowing the identification of cell type-specific cellular responses during fungal invasion. We further assigned infection site-associated cells using trajectory inference and live-cell imaging, which enabled the dissection of spatiotemporal dynamics in the cellular response during fungal colonisation. This study uncovered novel gene functions masked in studies using the bulk sequencing.

Our current knowledge of plant immunity is largely based on studies of the model bacterial pathogen *Pseudomonas syringae*, which colonises leaf apoplast. However, pathogens utilise diverse mechanisms to enter plant tissue and explore colonisation in different organs. Many pathogens can establish infection in the plant vascular system (Yadeta and J. Thomma, 2013). We observed that many genes encoding intracellular immune receptors (NLRs) were highly expressed in vasculature-related cells and in many cases, further induced by fungal infection in these specific cell types. As key components of plant immunity, NLRs recognise pathogen effectors delivered into plant cells and activate defence response. The specific expression of TNLs in procambium cells may indicate a role in defending xylem and phloem colonising pathogens, such as the bacterial pathogens *Xylella fastidiosa* (Yadeta and J. Thomma, 2013), phytoplasma (Sugio et al., 2011), *Ralstonia solanacearum* (Mori et al., 2016), and *Candidatus Liberibacter* spp. (Wang et al., 2017), as well as the fungal pathogens such as *Fusarium oxysporum* (Ma et al., 2010) and *Verticillium* spp. (Fradin and Thomma, 2006), and oomycete *Pythium ultimum* (Levesque et al., 2010). Most of these pathogens rely on effector proteins to suppress host immune systems. Although specific effector-NLR recognition in vasculature has been reported, for example, the tomato CNL I-2 recognizes the *F. oxysporum* effector Avr2 in the xylem sap (Takken and Rep, 2010), how NLR receptors, and indeed immune signalling, work in the vasculature tissues in general is poorly understood (Jiang et al., 2019). The fact that there is currently no known NLR-mediated resistance against the phloem-colonizing bacterial pathogens such as Liberibacter and phytoplasmas indicates that the classic ETI may not be deployed in mature vasculature tissues although these pathogens do secrete effectors (Clark et al., 2018; Sugio *et al*., 2011). Future research is warranted to understand the biological importance of the enriched expression of NLRs, especially TNLs, in the vasculature-related cells.

While NLRs are activated at the protein level by forming the resistosome protein complexes, some NLRs, particularly many of the TNLs, are induced at the transcript level after the activation of PTI (Bjornson et al., 2021; Tian et al., 2021) and ETI (Mine et al., 2018). Consistent with that, we found a family of TNLs to be highly induced in companion cells during an early infection stage of *C. higginsianum*, which is not known to invade vasculature. Therefore, this enhanced expression of TNLs likely represents a general response aiming to protect the vasculature organs, which are not only vital for plant growth and survival but could also be hijacked by pathogens for systematic infection. In contrast to TNLs, we did not observe a general cell type-specific expression pattern for CNLs, which may reflect a functional divergence between these two major classes of NLRs. Further investigations on this interesting phenomenon will offer novel insight into the cell type-specific functionality of plant NLRs.

A major challenge in the studies of plant-pathogen interactions is the uneven distribution of pathogen invasion/colonisation and, in the case of filamentous pathogens, the dynamic pathogen development in the host tissue (Giraldo and Valent, 2013; Latijnhouwers *et al*., 2003; Lo Presti et al., 2015). We determined the spatiotemporal heterogeneity in plant responses at the single cell resolution. Our analyses reveal a robust correlation between cellular response and the proximity of the cell to the invading fungal hyphae. A similar coupling of spatial and temporal responses was also observed during the bacterial infection (Zhu *et al*., 2022). More importantly, we uncovered a novel, cell-type-specific response that showed spatial dynamics. In this case, coordinated expression changes in both positive and negative regulators of ABA signalling occurred specifically in guard cells that had direct contact with fungal hyphae. This activation of ABA signalling gradually decreased in surrounding cells and diminished in more distal cells. Consistent with this transcription signature, the stomatal aperture gradually decreased in cells towards the fungal infection sites. Stomata formed by guard cells serve as natural entry points by bacterial pathogens (Lovelace and Ma, 2022; Melotto *et al*., 2006). It has been well established that stomatal closure is a defense response induced upon bacterial perception to prevent their entry to the apoplast (Hu *et al*., 2022; Roussin-Léveillée *et al*., 2022). Although filamentous pathogens can force their entry through mechanical pressures generated at appressoria (Ryder et al., 2022; Wilson and Talbot, 2009) or slicing through the epidermal cells (Bronkhorst et al., 2021; Bronkhorst et al., 2022), it has been observed that open stomata could also be exploited by oomycetes as an alternative entry mechanism and at the later infection stage for sporulation (Fawke *et al*., 2015; Wang et al., 2013). Further functional characterization is required to determine whether stomatal closure can contribute to preventing fungal/oomycete invasion and/or sporulation.

In addition to cell type-specific molecular events, our work also highlighted the importance of cell-type-specific gene activations in disease resistance. Indole glucosinolates (IG) are nitrogen- and sulphur-rich phytoanticipins that are critical to plant defence (Xu *et al*., 2016; Yang et al., 2020). The IG biosynthetic pathway is activated during immune signalling and the final products have been implicated in resistance against a variety of fungal and oomycete pathogens (Xu *et al*., 2016). Consistent with these previous findings, we observed an induction of IG biosynthesis in cells that had direct contact with *C. higginsianum*. Importantly, we found that IG production involved different genes in different cell types. In particular, MYB122, MYB34, and MYB51 were proposed to function redundantly in IG biosynthesis (Frerigmann and Gigolashvili, 2014). However, functional characterization demonstrated a significant contribution of MYB122 to plant defense, possibly through its specific regulation of IG production in epidermal cells at the infection site. This finding exemplifies the power of single cell-based analysis in dissecting gene functions.

This study provides a comprehensive single-cell atlas in Arabidopsis during the infection of a hemibiotrophic fungal pathogen. We show highly heterogenous gene expression profiles in plant cells representing the dynamic cellular response to a fungal infection that is not only affected by cell-type specificity but also relative proximity to fungal invasive growth. This high-resolution transcriptome dataset represents a rich resource that will facilitate studies to understand molecular details underlying plant-pathogen interactions. Genes and cis-elements are anticipated to be identified using this dataset, which will be useful to generate constructs that specifically express defence-related genes in the most relevant cells. In this manner, the effort of developing disease-resistant crops will be enhanced by minimising growth penalty.

## ACKNOWLEDGMENTS

We thank Professor Dan MacLean, Dr Clara Jégousse, and George Deeks for technical support on the bioinformatic tools and data management. We are grateful to Dr Christina Faulkner for kindly providing *C. higginsianum* strains. We are grateful to all members of the Ma group for helpful discussions and technical support. This project is supported by Gatsby Charitable Foundation and UKRI BBSRC Grant BBS/E/J/000PR9797.

## AUTHOR CONTRIBUTIONS

W.M. and B.T. conceived the research and designed the experiments. B.T. performed the experiments. B.T. and L.F. analysed the sequencing data. W.M., B.T. and P.D. wrote the manuscript.

## STAR METHODS

### CONTACT FOR REAGENT AND RESOURCE SHARING

Further information and requests for resources and reagents should be directed to and will be fulfilled by the Lead Contact, Wenbo Ma.

## DATA AND METERIAL AVAILABILITY

All code is available at https://github.com/tangbozeng/scrna_seq_fungal_infection. Materials are available from W.M. upon request under a materials transfer agreement with The Sainsbury Laboratory. The raw data of scRNA-seq and bulk RNA-seq were deposited in ENA with project ID PRJEB61052.

## METHOD DETAILS

### Plant material and growth conditions

*Arabidopsis thaliana* ecotype Col-0 (wild-type) seeds were sterilised and grown on Murashige-Skoog medium plates supplemented with 1% sucrose and 0.8% Phytagel in a growth chamber under 16/8-hour light/dark condition at 22°C. After 14 days, the seedlings were inoculated with conidia suspensions of *C. higginsianum* and used for scRNA-seq. For pathogenicity assays, Arabidopsis plants were grown in a growth room at 22°C with 16/8h light/dark regime.

### Inoculation with *C. higginsianum*

Growth of *C. higginsianum* was performed as previously described (O’Connell et al., 2004). *C. higginsianum* strain IMI 349061 was sub-cultured from −80 °C stock and grown in PDA plates at 25°C with a 12-hour light-dark cycle. After 5 days, conidia of *C. higginsianum* were harvested by being washed with water. The suspension was then used to make a conidial suspension containing 0.2% gelatin in concentration 1x 10^5^ conidia per mL. 14-day-old Arabidopsis seedlings were spray-inoculated with either the conidia suspension or 0.2% of gelatin as the mock control. The inoculated plants were kept in a growth chamber at 25°C with a 12-hour light-dark cycle. Arabidopsis mutant lines of *myb122* were ordered from salk lines (Salk_027525 and Salk_27085) and verified by PCR and qRT-PCR. The primers used to genotype and test the expression of these mutants are listed in the key resources table. Four-week-old Arabidopsis plants were inoculated with the conidial suspension, and 20 µL conidial suspension was applied to the abaxial side of each adult rosette leaves. The plants were kept with high humidity in a growth chamber under a 12-hour light-dark cycle for five days. Inoculated leaves were collected and immediately scanned. The discoloured area (DA) and total area (TA) of each leaf were determined by the Image-Adjust-ColorThreshold functions of the ImageJ-fuji software. The lesion size is represented by DA/TA.

### Transient expression of MYB122 in *Nicotiana benthamiana*

To validate the specific induction of *MYB122* in epidermal cells at the site of fungal infection, *pMYB122:MYB122-YFP* was constructed by cloning a 1.5 kb DNA sequence upstream of the *MYB122-* encoding gene with the cDNA sequence of *MYB122* fused with *YFP* and then was introduced into vector PGWB514. The constructed plasmid carrying *pMYB122:MYB122-YFP* was introduced into *Agrobacterirum tumefaciens*, which was then used to infiltrate into leaves of *N. bethaminana*. At 24 hours post Agro-infiltration, the leaves were inoculated with the conidial suspension of *C. higginsianum* that constitutively expressed GFP. At 36 hours post fungal inoculation, leaves were collected for imaging analysis.

### Microscope for live-cell imaging and stomatal aperture quantification

The inoculated leaf was all collected from seedlings tissue after infection unless specified. The fluorescence signal was observed under a Leica TCL SP8 confocal microscope. A laser of 488-nm wavelength was used for excitation, and an emission wavelength of 500-530 nm was used to observe the GFP signal constitutively expressed in *C. higginsianum*. To acquire the Venus signal, a 514-nm laser was used for excitation, and the emission detection wavelength was between 515 nm and 545 nm. To measure stomatal aperture, leaves from 14-day seedlings were used by being treated with water with 0.1% gelatine and suspension of fungal spores as mock and infected samples. The samples were left in a controlled environment under 16/8-hour light/dark condition at 22°C. At 40 hpi, the samples were collected for imaging acquisition. The image files capturing the stomatal aperture of uninfected, distant, or proximity guard cells were fed into a machine learning-based analysis pipeline (Sai *et al*., 2022). The width and length of each guard cell were measured and collected for comparison.

### Arabidopsis protoplast preparation for scRNA-seq

Before sample collection, live-cell imaging was performed to monitor the robustness of the infection, and ensure the infection was appreciated as expected. The protoplast isolation from plant leaves was mainly processed as described with minor modification (Yoo et al., 2007). Briefly, the mock and infected leaves were cut into 0.5 mm strips and immediately transferred into protoplast enzyme solution (4% cellulase R10, 1.5% macerozyme R10, 0.4 M mannitol, 10 mM KCl, 10 mM CaCl_2_, and 0.1% BSA) through 0.2 mm filter. At least 20 leaves of each sample were collected and bulked for further experiment. To shorten the digestion time, the leaf strips were wrapped in foil to avoid light and gently infiltrating in the vacuum for 20 min, and these enabled cells released from seedlings faster than usual. After 1h of digestion at room temperature, the leaves were fully digested. The protoplast was filtered 3-4 via cell strainers and washed 2 times with pre-cooled 10% PBS buffer at 4°C in a 50 mL sterile tube. The samples were centrifuged at 150 x g for 2 min at 4°C and then transferred into ice. The suspension was kept on ice for at least 10 min to allow the protoplasts to fall down onto the bottom of the tube so that the upper suspension could be discarded without disturbing the protoplasts. The viability of protoplasts and the cleanness of the samples was determined by trypan blue staining. The samples with >90% viability were adjusted to ∼2000 cells / µL using a hemocytometer and proceeded for single-cell sorting. At least four technical replicates for each biological replicate were prepared, and the best two were selected for further single-cell partition.

### Single-cell partition and RNA-library construction

The protoplast suspensions were immediately loaded into a Chromium Single Cell Instrument (10x Genomics, Pleasanton, CA) to produce single-cell GEMs (Gel Bead-In Emulsions). scRNA-seq libraries were generated using the Chromium Single Cell 3’ Gel Bead and Library Kit V3.1 (10x Genomics, Pleasanton, CA). The experiment was performed according to the user’s guide provided by the vendor. The libraries were sequenced by Illumina NovaSeq (Novo Gene, Cambridge, UK) after quality control by Agilent 2100 Bioanalyzer.

### Alignment of raw scRNA-seq data

The reference files for Arabidopsis genome were acquired and downloaded from Ensembl database (Cunningham et al., 2022). The “cellranger mkref” function was used to build references. Raw reads were alignment against the Araport11 using Cellranger (V6.0.1, 10x Genomics, Pleasanton, CA) with default settings. The percentage of aligned reads was ranged from 70% to 95% in the samples.

### Quality control, doublet detection, cell-cycle regression, and batch effects removal of scRNA-seq data

The downstream analysis was conducted by deploying scripts modified from the Seurat (v4.0) (Hao et al., 2021). Read10x was used to load raw matrix files, and the function “CreateSeuratObject” was used to build individual Seurat dataset after Cellranger alignment. To filter out the low-quality data, the cells with less than 200 genes were discarded, and the genes present less than 3 cells were not considered. The dataset was then normalised by “SCTransform” function and fed into Douletfinder workflow to identify doublets during the single-cell instrument (McGinnis et al., 2019). Briefly, the function “DoubletFinder_v3” was used to determine homotypic doublets with parameters of nExp = round (0.05*nrow(x)) (the number of expected real doublets), pN=0.25 (the number of artificial doublets), and pK =0.09 (the neighbourhood size). The proportion of artificial neighbours for each cell was calculated as thereby used to define doublet predictions. The resultant cells annotated as “Singlets” in each library were kept for further analysis. The proportion of UMI aligned to chloroplast, mitochondrial, and produced-induce genes in each library were investigated and visualised before further analysis. The cells with no more than 10% of mitochondrial or chloroplast UMI were kept. Further quality control was then performed, as low-quality cells were defined as expressing less than 500 genes or more than 50,000 genes. The libraries were then merged by Seurat. To mitigate the effects of cell cycle heterogenicity on cell clustering, the genes related to G1-S, and G2-M were pulled out from the public database (Berardini et al., 2015) and fed with in the function “CellCycleScoring”. For normalisation, the SCTransform was applied but with “var.to.regress” for cell cycle genes regression. The resultant dataset was fed into “RunHarmony” function using seurat dataset assay as “SCT” to correct batch effects between the replicates and samples and generated integrated Seurat dataset (Korsunsky et al., 2019).

### Clustering and cell type annotation and gene expression analysis in scRNA-seq

Cluster analysis was performed by RunUMAP and RunTSNE following the clusters identification based on Louvain by inputting was constructed by “FindNeighbors” with parameters of k.param=10, dims=1:30, and “FindClusters” with resolution 0.4. A correlation analysis using all the libraries were performed using aggregated expression values from all the cells, and the libraries generated from mock at 24 hpi and 40 hpi were merged, as higher correlation values was yielded than that comparing to infected samples. To define cluster-enriched genes in the atlas, “FindAllMarkers” function was used with parameters of logfc.threshold = 0.4, min.pct=0.1, min.diff.pct=0.1. The Wilcoxon Rank Sum test method was used to define the differentially expressed genes between the two groups of cells. The resultant genes were then merged with single-cell marker genes database (Chen *et al*., 2021; Xu *et al*., 2022) and the supplementation data collected from previous studies using Arabidopsis leaf tissue (Kim *et al*., 2021).The resultant genes were used for cell type annotation of each cluster. To visualize and profile marker gene expression, Featureplot function was applied. For sub-clustering analysis of cluster 6 and 18, the cells were subset and processed by RunUMAP with resolution 0.8 to yield higher number of clusters. The sub-clusters enriched genes and cell type annotations were performed as described above. To define relative expression of putative and known NLR-encoding genes in Arabidopsis, we collected the data generated from previous study using NLRtracker pipeline (Kourelis *et al*., 2021), Only genes expressed in at least 500 cells were kept for further investigation. The UMI count matrix generated from Seurat was then pull out, and the relative expression of each gene in each cell type was then derived by creating a matrix with aggregated expression values for defined groups of cells after being divided by cell size factors (Cao *et al*., 2019). To investigate the expression of pattern of CNL, TNL, and RNL gene family, the NLR-encoding genes for each sub-family were pooled together as input for AddModuleScore function in Seurat (Hao *et al*., 2021). This method defined relative contribution of the targeted gene families to each cell. Here, we deployed scripts with default setting “ctrl = 50” as 50 randomly picked genes as housekeeping gene were used for benchmarking.

### Trajectory inference and pseudotime analysis

To construct a trajectory, cells annotated with the same cell type were pooled together and fed into a monocle pipeline (Trapnell et al., 2014). To do this, the UMI count matrix for the four major cell types as ‘single.cell.experiment’ was used to construct the dataset by “newCellDataSet” function, respectively. The resultant dataset was processed to identify genes that were differentially expressed while eliminating batch effects from replicates. The top 2,000 significantly altered genes (according to the q-values) from this dataset were selected for further analysis. To minimise the impact of cell type developmental effects on infection-related trajectory construction, uninfected cells were isolated from each dataset, and wild-type pseudotime was inferred to identify the top 100 significantly altered genes that are crucial for cell type development and determinant. The first set of genes, excluding those identified in the cell-type pseudotime analysis, was selected to construct the infection-responsive trajectory, which is vital to build the trajectory elucidating the dynamics of cellular responses to infection. To build trajectory, setOrderingFilter was applied and DDRTree method was used to order the cells according to the expression pattern of selected genes. The resultant pseudotime values for four cell types were then yielded to illustrate infection progression. To define the expression of ABA-related genes, genes encoding ABA positive regulators and negative regulators were pulled out from ATIR database (Berardini *et al*., 2015), respectively. The resultant gene lists were input into AddModuleScore, and relative expression of ABA-related genes in each cell was yielded for further investigation. To define genes enriched at infection sites, the cells with the top 10% pseudotime defined from the trajectory of each cell type was pulled out and defined to serve as infection sites. After that the FindAllMarkers function in Seurat was applied to defined genes enriched in the cells at infection sites by settings with Log2|FC|>0.25, pct.1>=0.2, pct.2<0.1, min.diff.pct>0.2, using The Wilcoxon Rank Sum test method. All the enrichment analysis was performed using clusterProfiler package (Wu et al., 2021) with threshold of q-value<0.2 as cut-off for KEGG metabolic, and q-value<0.2 for GO terms.

### Total RNA isolation and Bulked RNA-seq analysis

To determine protoplasting-induced genes, bulked RNA-seq was performed. Leaves from 14-day old seedlings were cut into 0.5 mm strips and digested in protoplast enzyme solution as described above. The protoplasts were collected and processed for total RNA isolation by Qiagen kit (key resource table) according to instructions provided by the manufacturer. Leaves directly collected from the seedlings without treatment were used as a mock control. The RNA extracts were treated with RNase-Free DNAase. The elimination of genome DNA was checked by PCR using primers amplifying an intron region of the actin-encoding gene in Arabidopsis. For bulked RNA-seq to determine transcriptional profiling of Arabidopsis during infection by *C. higginsianum*, the raw data was downloaded from NCBI datasets from ENA under projects PRJNA148307 and PRJNA151285 (O’Connell *et al*., 2012). The raw reads of all the bulked RNA-seq were fed for quality check and adaptor removal by Trimmomatic (Bolger and Giorgi, 2014). Clean reads were aligned against Arabidopsis genome using Kallisto (Bray et al., 2016) followed by tximport (Soneson et al., 2015) to generate a count matrix. Batch effects removal was performed by sva package (Leek et al., 2012). Sample size factors, normalisation, and differentially expressed genes were analysed by DESeq2 (Love et al., 2014). To identify Arabidopsis genes specifically induced during fungal infection, the genes showing log_2_FC >2, p-adj<0.05, and averaged TPM <100 in the mock sample were considered as appressorium stage induced genes as an early response, and the ones at 42 hpi were considered as biotrophic stage-specific.

---

**Figure S1.**
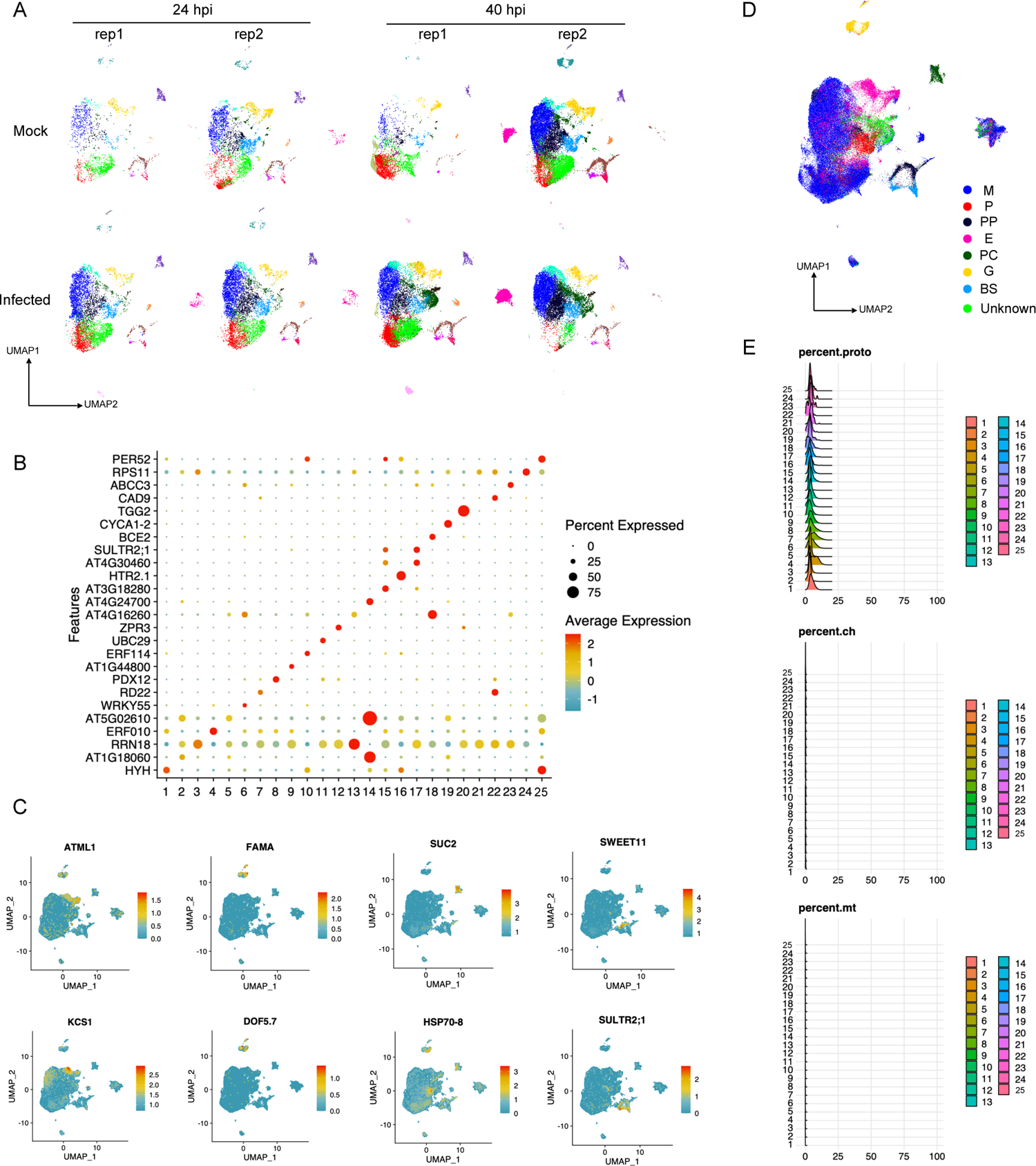
Construction of an Arabidopsis leaf cell transcriptome atlas, related to Figure 1. (**A**) UMAP visualisation of single cell atlas showing cell clustering from each replicate of all the samples. Each dot represents the transcriptional signature of a single cell. A total of 25 clusters are represented by different colours. (**B**) Representative genes enriched in each cluster that were used for assigning the cell type of each cluster. The dotplot shows the specific expression of the cluster-enriched genes in each cell cluster. The colour of the circles indicates relative transcript abundance and the size of the circles indicates the percentage of cells expressing the specific gene. This analysis was done by Wilcoxon Rank Sum Test, which enabled to set cut-off as 1.5-fold changes, and 20% of cells showing higher transcripts than all other cells in the atlas.(**C**) Expression profiling of cell-type marker genes including *ATML1* (Epidermis), *KCS1* (Epidermis), *FAMA* (guard cell), *DOF5.7* (guard cell), *SUC2* (phloem companion), *HSP70-8* (Procambium), *SULTR2;1* (phloem companion), and *SWEET11* (phloem parenochyma) in the scRNA-seq atlas visualized by UMAP. Colours indicate relative expression. (**D**) Single-cell atlas showing the cell type assignment. (**E**) Ridge-plots showing the transcript fraction of protoplasting-induced genes (top), chloroplast genes (middle), and mitochondrional genes (bottom) in each cluster. X-axis shows proportion of the gene transcripts in each cell, and the y-axis shows the relative number of cells.

**Figure S2.**
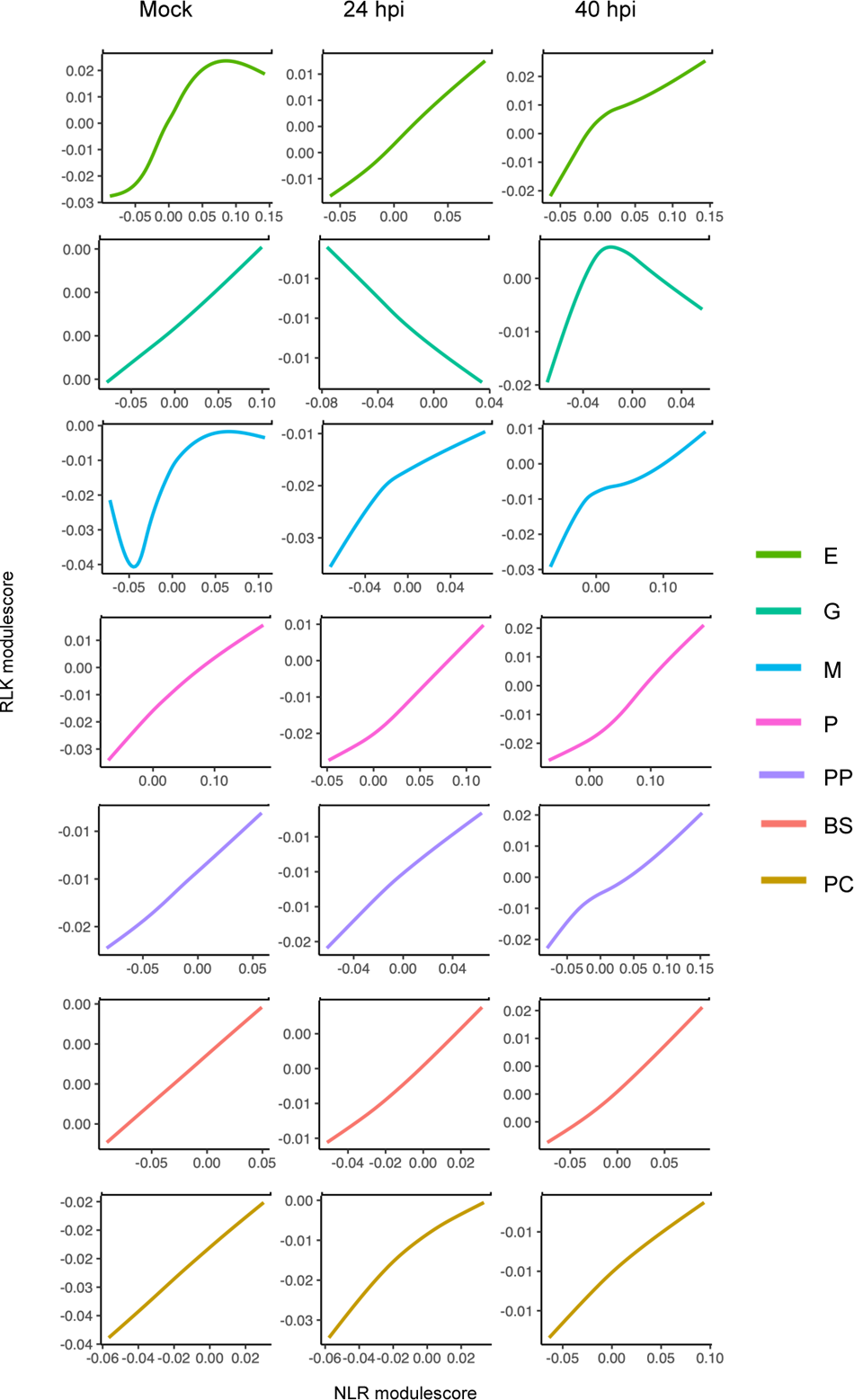
Correlated expression of genes encoding immune receptors NLRs and RLKs, related to Figure 2. The linear regression line plots show correlation of relative expression of *NLR*s (X-axis) and *RLK*s (Y-axis) as module-scores in cells. Only cells with cell type annotation were analyzed. Putative NLR or RLK encoding genes were either predicted by NLRtracker (Kourelis *et al*., 2021), or pulled out from the PRGdb 3.0 database (Osuna-Cruz et al., 2018), respectively.

**Figure S3.**
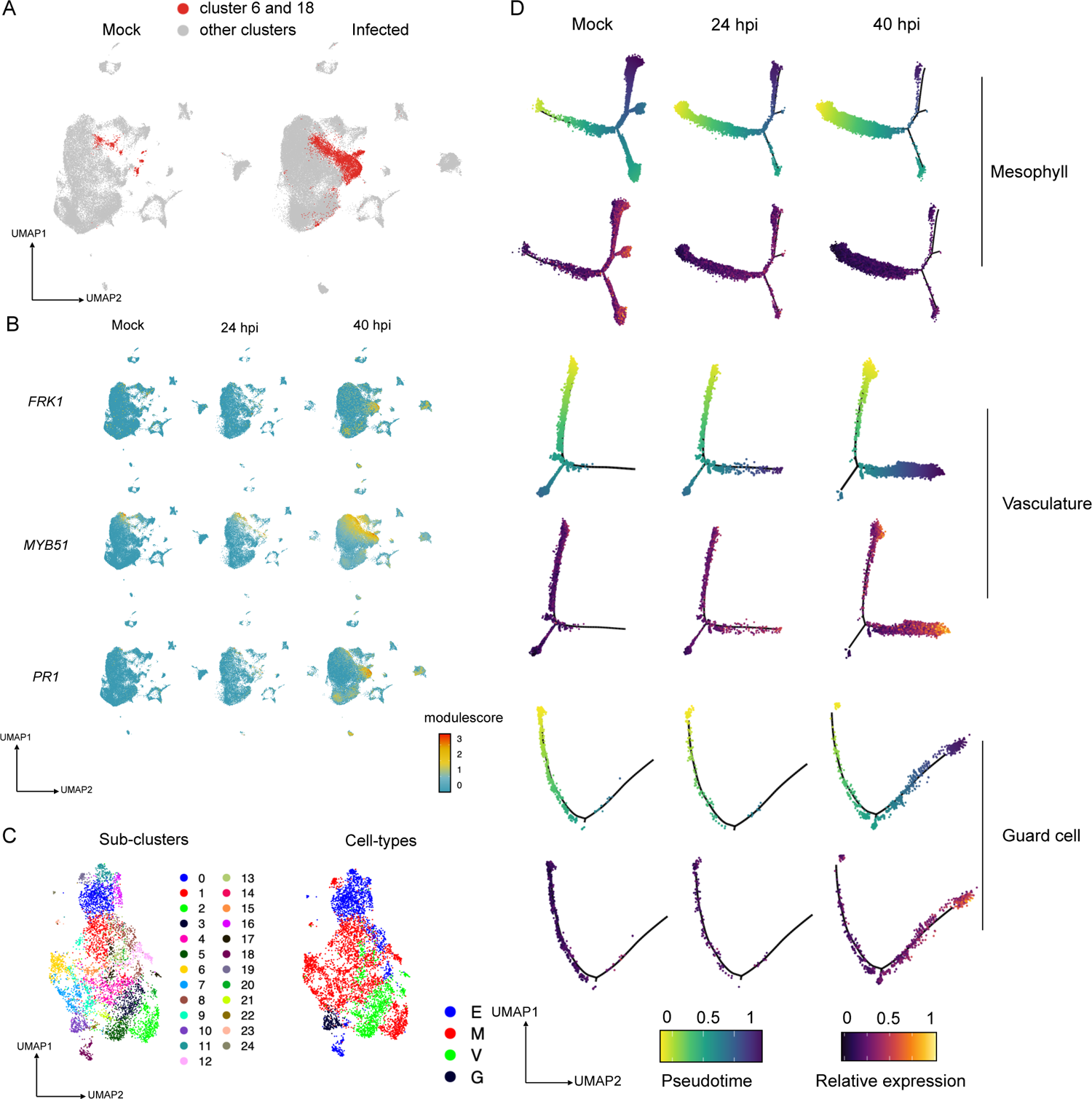
Profiling of infection-related cell clusters and their sub-clustering, related to Figure 3. (**A**) Cells belonging to the clusters 6 and 18 were over-representative in infected samples. These two clusters could not be properly assigned to a cell type when being analyzed together the total cell population. Cells constituting cluster 6 and 18 are highlighted as red dots in the atlas. (**B**) Transcripts of genes involved in defense response were enriched in cluster 6 and 18 cells. The relative expression of three immunity-related markers, *FRK1*, *MYB51*, and *PR1*, are presented in each cell in the atlas. (**C**) Sub-clustering enabled cell type annotation for the cells from clusters 6 and 18. Left panel: An atlas built on cells from the clusters 6 and 18 with the colours representing sub-clustering. Right panel: the cell atlas with cell type annotation with the colours representing the four major cell types based on expression of cell type marker genes. Vasculature cells are not further annotated due to the smaller cell number. (**D**) A shift of cell population in response to fungal infection observed in each of the four cell types. Upper panel: trajectory curves of mesophyll, vasculature, and guard cells from mock, 24 hpi and 40 hpi samples with each dot representing a cell. Colours of the dots represent their pseudotime values. Lower panel: aggregated expression level of a set of 41 genes that were identified to be induced at the infection site of powdery mildew (Chandran *et al*., 2010) mapped to the trajectory curves. Colour intensity indicates the relative value of module-score in each cell.

**Figure S4.**
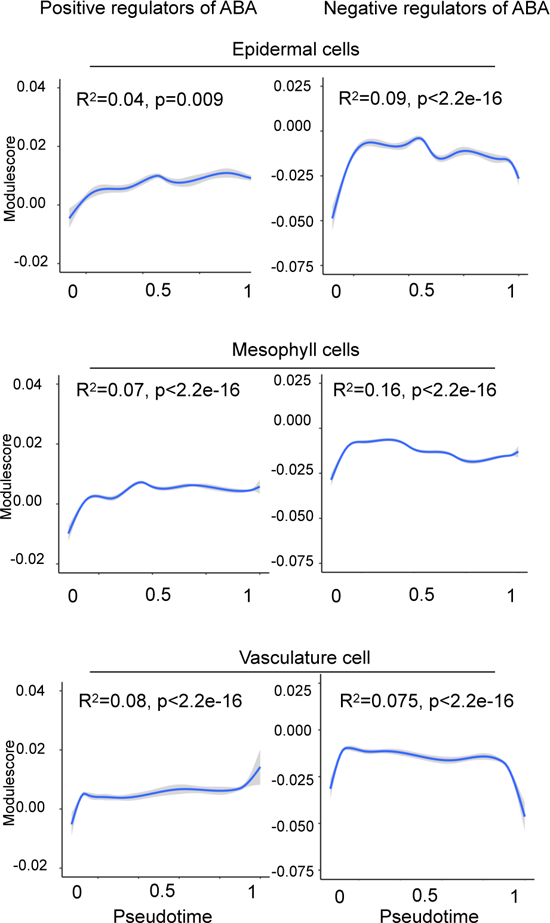
Genes encoding ABA-related regulators did not show expression changes along the trajectory curves in epidermal, mesophyll, and vasculature cells, related to Figure 4. Linear regression line plots show expression patterns of genes encoding positive (left) or negative (right) regulators of ABA signalling. The X-axis represents pseudotime values assigned to the cells in the infection-responsive trajectory and the Y-axis represents relative expression levels as module-scores.

**Figure S5.**
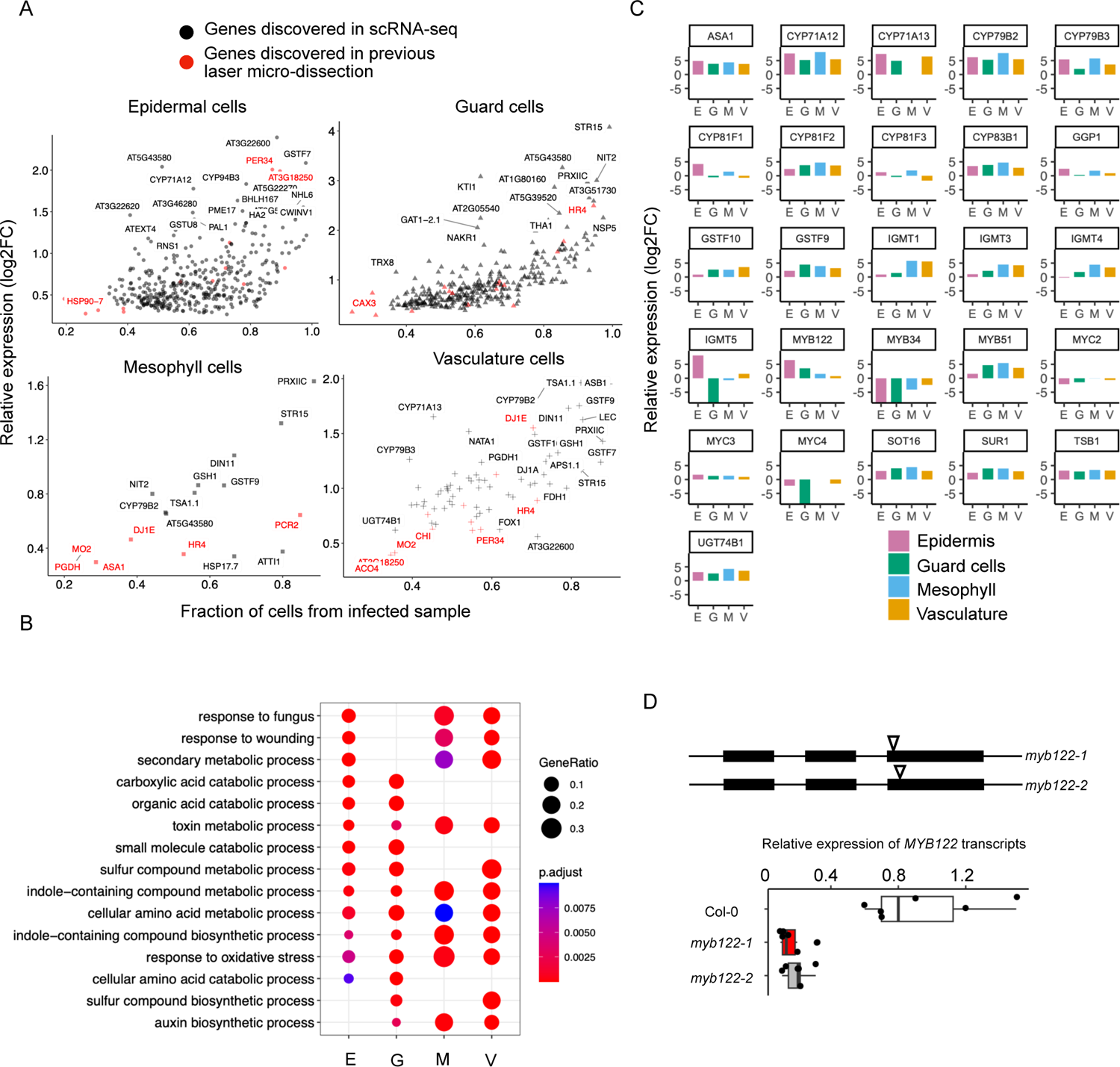
Discovery of genes exhibiting cell type-specific expression at the infection sites, related to figure 5. (**A**) scRNA-seq analysis discovered many genes with induced expression in cells at the infection site and with a cell-type-specific manner. The dot-plots show the relative expression of genes significantly induced at infection site in each of the four cell types. The X-axis represents the percentage of the cells with the corresponding gene induced. The Y-axis represents the log_2_FC value of each gene in cells at the infection sites vs all other cells using the whole scRNA-seq dataset. Genes in red were identified as associated with infection sites of powdery mildew from a laser microdissection study. (**B**) Biological processes represented by GO terms enriched in infection sites in the four cell types. Dot-plot was generated using genes transcriptionally induced at infection sites. (**C**) Genes encoding components involved in IGs biosynthesis exhibited plasticity in cell type-specific expression at infection sites. The plots show the relative expression level (log2|FC|) of the genes in infection sites from each of the four cell types. The values were generated by comparing the relative transcript abundance of each gene in the infection site cell in a specific cell type to that in all other cells in the same cell type. (**D**) Confirmation of *MYB122* mutant of Arabidopsis. Two independent T-DNA insertion lines, *my122-1* (Salk_027525) and *myb122-2* (Salk_27085), were examined. The schematic plot shows the T-DNA insertion sites of the mutants, which were confirmed by PCR and sequencing. The lack of *MYB122* expression was also confirmed by quantitative RT-PCR. Genes encoding actin and beta-tubulin were used as internal controls for the qRT-PCR analysis.

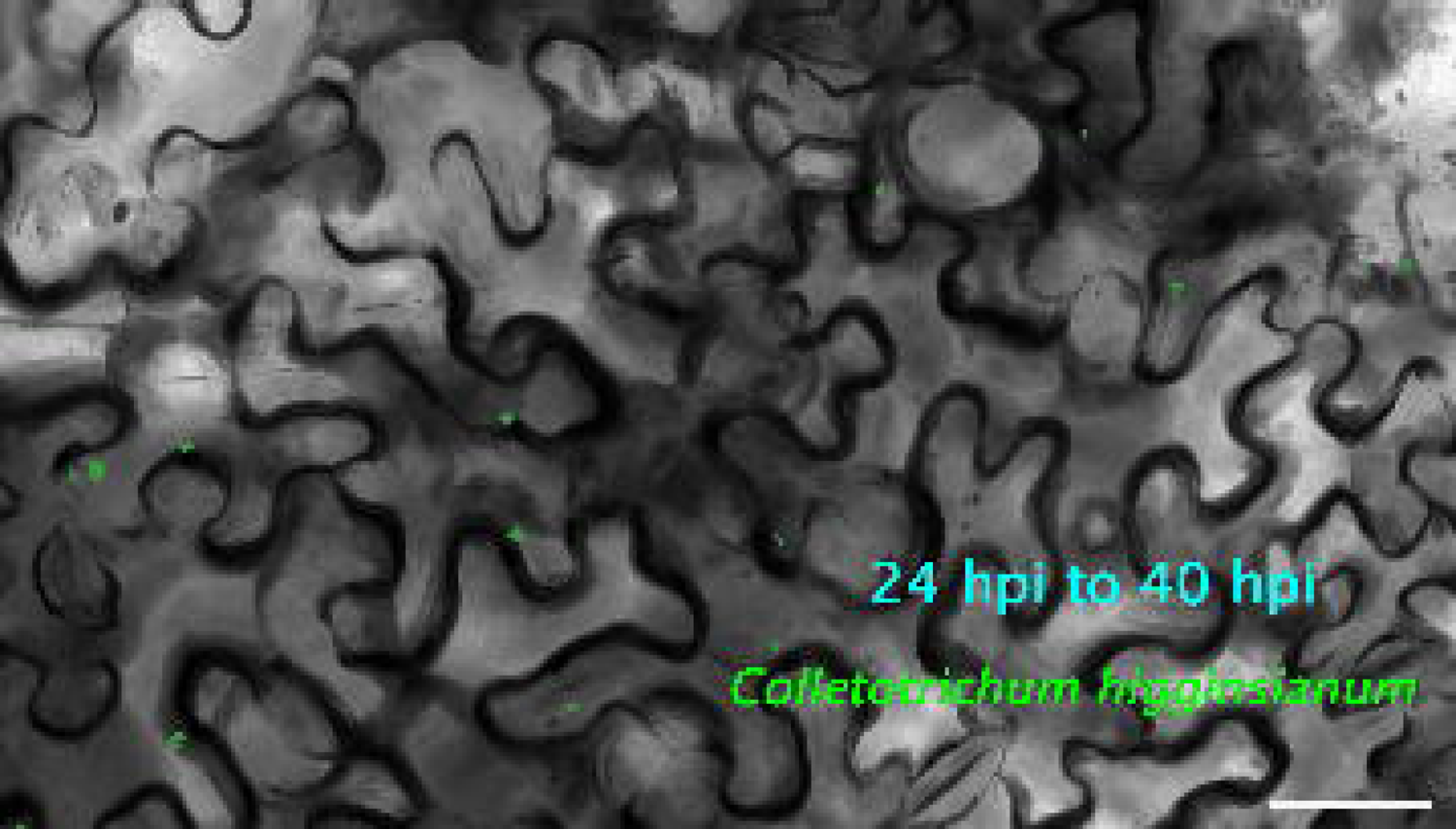

## REFERENCES

Aerts, N., Chhillar, H., Ding, P., and Van Wees, S.C. (2022). Transcriptional regulation of plant innate immunity. Essays in Biochemistry 66, 607–620.

Badel, J.L., Shimizu, R., Oh, H.-S., and Collmer, A. (2006). A Pseudomonas syringae pv. tomato avrE1/hopM1 mutant is severely reduced in growth and lesion formation in tomato. Molecular plant-microbe interactions 19, 99–111.

Batool, S., Uslu, V.V., Rajab, H., Ahmad, N., Waadt, R., Geiger, D., Malagoli, M., Xiang, C.-B., Hedrich, R., and Rennenberg, H. (2018). Sulfate is incorporated into cysteine to trigger ABA production and stomatal closure. The Plant Cell 30, 2973–2987.

Berardini, T.Z., Reiser, L., Li, D., Mezheritsky, Y., Muller, R., Strait, E., and Huala, E. (2015). The Arabidopsis information resource: making and mining the “gold standard” annotated reference plant genome. genesis 53, 474–485.

Birkenbihl, R.P., Liu, S., and Somssich, I.E. (2017). Transcriptional events defining plant immune responses. Current Opinion in Plant Biology 38, 1–9.

Bjornson, M., Pimprikar, P., Nürnberger, T., and Zipfel, C. (2021). The transcriptional landscape of Arabidopsis thaliana pattern-triggered immunity. Nature plants 7, 579–586.

Bolger, A., and Giorgi, F. (2014). Trimmomatic: A flexible read trimming tool for Illumina NGS data. Bioinformatics 30, 2114–2120.

Bozkurt, T.O., and Kamoun, S. (2020). The plant–pathogen haustorial interface at a glance. Journal of Cell Science 133, jcs237958.

Bray, N.L., Pimentel, H., Melsted, P., and Pachter, L. (2016). Near-optimal probabilistic RNA-seq quantification. Nature biotechnology 34, 525–527.

Bronkhorst, J., Kasteel, M., van Veen, S., Clough, J.M., Kots, K., Buijs, J., van der Gucht, J., Ketelaar, T., Govers, F., and Sprakel, J. (2021). A slicing mechanism facilitates host entry by plant-pathogenic Phytophthora. Nature Microbiology 6, 1000–1006.

Bronkhorst, J., Kots, K., de Jong, D., Kasteel, M., van Boxmeer, T., Joemmanbaks, T., Govers, F., van der Gucht, J., Ketelaar, T., and Sprakel, J. (2022). An actin mechanostat ensures hyphal tip sharpness in Phytophthora infestans to achieve host penetration. Science Advances 8, eabo0875.

Cao, J., Spielmann, M., Qiu, X., Huang, X., Ibrahim, D.M., Hill, A.J., Zhang, F., Mundlos, S., Christiansen, L., and Steemers, F.J. (2019). The single-cell transcriptional landscape of mammalian organogenesis. Nature 566, 496–502.

Cervantes-Pérez, S.A., Thibivilliers, S., Laffont, C., Farmer, A.D., Frugier, F., and Libault, M. (2022). Cell-specific pathways recruited for symbiotic nodulation in the Medicago truncatula legume. Molecular Plant 15, 1868–1888.

Chandran, D., Inada, N., Hather, G., Kleindt, C.K., and Wildermuth, M.C. (2010). Laser microdissection of Arabidopsis cells at the powdery mildew infection site reveals site-specific processes and regulators. Proceedings of the National Academy of Sciences 107, 460–465.

Chen, H., Yin, X., Guo, L., Yao, J., Ding, Y., Xu, X., Liu, L., Zhu, Q.-H., Chu, Q., and Fan, L. (2021). PlantscRNAdb: a database for plant single-cell RNA analysis. Molecular Plant 14, 855–857.

Clark, K., Franco, J.Y., Schwizer, S., Pang, Z., Hawara, E., Liebrand, T.W., Pagliaccia, D., Zeng, L., Gurung, F.B., and Wang, P. (2018). An effector from the Huanglongbing-associated pathogen targets citrus proteases. Nature communications 9, 1718.

Cui, H., Tsuda, K., and Parker, J.E. (2015). Effector-triggered immunity: from pathogen perception to robust defense. Annual review of plant biology 66, 487–511.

Cunningham, F., Allen, J.E., Allen, J., Alvarez-Jarreta, J., Amode, M.R., Armean, I.M., Austine-Orimoloye, O., Azov, A.G., Barnes, I., and Bennett, R. (2022). Ensembl 2022. Nucleic acids research 50, D988–D995.

De Rybel, B., Mähönen, A.P., Helariutta, Y., and Weijers, D. (2016). Plant vascular development: from early specification to differentiation. Nature reviews Molecular cell biology 17, 30–40.

De Silva, D.D., Crous, P.W., Ades, P.K., Hyde, K.D., and Taylor, P.W. (2017). Life styles of Colletotrichum species and implications for plant biosecurity. Fungal Biology Reviews 31, 155–168.

Dodds, P.N., and Rathjen, J.P. (2010). Plant immunity: towards an integrated view of plant–pathogen interactions. Nature Reviews Genetics 11, 539-548.

Fawke, S., Doumane, M., and Schornack, S. (2015). Oomycete interactions with plants: infection strategies and resistance principles. Microbiology and Molecular Biology Reviews 79, 263–280.

Fradin, E.F., and Thomma, B.P. (2006). Physiology and molecular aspects of Verticillium wilt diseases caused by V. dahliae and V. albo-atrum. Molecular plant pathology 7, 71-86.

Frerigmann, H., and Gigolashvili, T. (2014). MYB34, MYB51, and MYB122 distinctly regulate indolic glucosinolate biosynthesis in Arabidopsis thaliana. Molecular Plant 7, 814-828.

Giraldo, M.C., and Valent, B. (2013). Filamentous plant pathogen effectors in action. Nature Reviews Microbiology 11, 800–814.

Gong, Y., Tian, L., Kontos, I., Li, J., and Li, X. (2023). Plant immune signaling network mediated by helper NLRs. Current Opinion in Plant Biology 73, 102354. https://doi.org/10.1016/j.pbi.2023.102354.

Hao, Y., Hao, S., Andersen-Nissen, E., Mauck III, W.M., Zheng, S., Butler, A., Lee, M.J., Wilk, A.J., Darby, C., and Zager, M. (2021). Integrated analysis of multimodal single-cell data. Cell 184, 3573–3587. e3529.

Hu, Y., Ding, Y., Cai, B., Qin, X., Wu, J., Yuan, M., Wan, S., Zhao, Y., and Xin, X.-F. (2022). Bacterial effectors manipulate plant abscisic acid signaling for creation of an aqueous apoplast. Cell Host & Microbe 30, 518–529. e516.

Illouz-Eliaz, N., Lande, K., Yu, J., Jow, B., Swift, J., Lee, T., Nobori, T., Castanon, R.G., Nery, J.R., and Ecker, J.R. (2023). Drought Recovery Induced Immunity Confers Pathogen Resistance. bioRxiv, 2023.2002. 2027.530256.

Jiang, Y., Zhang, C.-X., Chen, R., and He, S.Y. (2019). Challenging battles of plants with phloem-feeding insects and prokaryotic pathogens. Proceedings of the National Academy of Sciences 116, 23390–23397.

Jouannet, V., Brackmann, K., and Greb, T. (2015). (Pro) cambium formation and proliferation: two sides of the same coin? Current opinion in plant biology 23, 54–60.

Khang, C.H., Berruyer, R., Giraldo, M.C., Kankanala, P., Park, S.-Y., Czymmek, K., Kang, S., and Valent, B. (2010). Translocation of Magnaporthe oryzae effectors into rice cells and their subsequent cell-to-cell movement. The Plant Cell 22, 1388–1403.

Kim, J.-Y., Symeonidi, E., Pang, T.Y., Denyer, T., Weidauer, D., Bezrutczyk, M., Miras, M., Zöllner, N., Hartwig, T., and Wudick, M.M. (2021). Distinct identities of leaf phloem cells revealed by single cell transcriptomics. The Plant Cell 33, 511–530.

Koeck, M., Hardham, A.R., and Dodds, P.N. (2011). The role of effectors of biotrophic and hemibiotrophic fungi in infection. Cellular microbiology 13, 1849–1857.

Korsunsky, I., Millard, N., Fan, J., Slowikowski, K., Zhang, F., Wei, K., Baglaenko, Y., Brenner, M., Loh, P.-r., and Raychaudhuri, S. (2019). Fast, sensitive and accurate integration of single-cell data with Harmony. Nature methods 16, 1289–1296.

Kourelis, J., Sakai, T., Adachi, H., and Kamoun, S. (2021). RefPlantNLR is a comprehensive collection of experimentally validated plant disease resistance proteins from the NLR family. PLoS Biology 19, e3001124.

Latijnhouwers, M., de Wit, P.J., and Govers, F. (2003). Oomycetes and fungi: similar weaponry to attack plants. Trends in microbiology 11, 462–469.

Leek, J.T., Johnson, W.E., Parker, H.S., Jaffe, A.E., and Storey, J.D. (2012). The sva package for removing batch effects and other unwanted variation in high-throughput experiments. Bioinformatics 28, 882–883.

Levesque, C.A., Brouwer, H., Cano, L., Hamilton, J.P., Holt, C., Huitema, E., Raffaele, S., Robideau, G.P., Thines, M., and Win, J. (2010). Genome sequence of the necrotrophic plant pathogen Pythium ultimum reveals original pathogenicity mechanisms and effector repertoire. Genome biology 11, 1–22.

Lindow, S.E., and Brandl, M.T. (2003). Microbiology of the phyllosphere. Applied and environmental microbiology 69, 1875–1883.

Liu, Z., Zhou, Y., Guo, J., Li, J., Tian, Z., Zhu, Z., Wang, J., Wu, R., Zhang, B., and Hu, Y. (2020). Global dynamic molecular profiling of stomatal lineage cell development by single-cell RNA sequencing. Molecular plant 13, 1178–1193.

Lo Presti, L., Lanver, D., Schweizer, G., Tanaka, S., Liang, L., Tollot, M., Zuccaro, A., Reissmann, S., and Kahmann, R. (2015). Fungal effectors and plant susceptibility. Annual review of plant biology 66, 513–545.

Lopez-Anido, C.B., Vatén, A., Smoot, N.K., Sharma, N., Guo, V., Gong, Y., Gil, M.X.A., Weimer, A.K., and Bergmann, D.C. (2021). Single-cell resolution of lineage trajectories in the Arabidopsis stomatal lineage and developing leaf. Developmental cell 56, 1043–1055. e1044.

Love, M.I., Huber, W., and Anders, S. (2014). Moderated estimation of fold change and dispersion for RNA-seq data with DESeq2. Genome biology 15, 1–21.

Lovelace, A.H., and Ma, W. (2022). How do bacteria transform plants into their oasis? Cell Host & Microbe 30, 412–414.

Ma, L.-J., Van Der Does, H.C., Borkovich, K.A., Coleman, J.J., Daboussi, M.-J., Di Pietro, A., Dufresne, M., Freitag, M., Grabherr, M., and Henrissat, B. (2010). Comparative genomics reveals mobile pathogenicity chromosomes in Fusarium. Nature 464, 367–373.

Mazutis, L., Gilbert, J., Ung, W.L., Weitz, D.A., Griffiths, A.D., and Heyman, J.A. (2013). Single-cell analysis and sorting using droplet-based microfluidics. Nature protocols 8, 870–891.

McDowell, J.M. (2013). Genomic and transcriptomic insights into lifestyle transitions of a hemi-biotrophic fungal pathogen. New Phytologist 197, 1032–1034.

McGinnis, C.S., Murrow, L.M., and Gartner, Z.J. (2019). DoubletFinder: doublet detection in single-cell RNA sequencing data using artificial nearest neighbors. Cell systems 8, 329–337. e324.

McLachlan, D.H., Kopischke, M., and Robatzek, S. (2014). Gate control: guard cell regulation by microbial stress. New Phytologist 203, 1049–1063.

Melotto, M., Underwood, W., Koczan, J., Nomura, K., and He, S.Y. (2006). Plant stomata function in innate immunity against bacterial invasion. Cell 126, 969–980.

Mine, A., Seyfferth, C., Kracher, B., Berens, M.L., Becker, D., and Tsuda, K. (2018). The defense phytohormone signaling network enables rapid, high-amplitude transcriptional reprogramming during effector-triggered immunity. The Plant Cell 30, 1199–1219.

Mori, Y., Inoue, K., Ikeda, K., Nakayashiki, H., Higashimoto, C., Ohnishi, K., Kiba, A., and Hikichi, Y. (2016). The vascular plant−pathogenic bacterium R alstonia solanacearum produces biofilms required for its virulence on the surfaces of tomato cells adjacent to intercellular spaces. Molecular Plant Pathology 17, 890–902.

Münch, S., Lingner, U., Floss, D.S., Ludwig, N., Sauer, N., and Deising, H.B. (2008). The hemibiotrophic lifestyle of Colletotrichum species. Journal of plant physiology 165, 41–51.

Ngou, B.P.M., Ding, P., and Jones, J.D. (2022). Thirty years of resistance: Zig-zag through the plant immune system. The Plant Cell 34, 1447–1478.

Nobori, T., Velásquez, A.C., Wu, J., Kvitko, B.H., Kremer, J.M., Wang, Y., He, S.Y., and Tsuda, K. (2018). Transcriptome landscape of a bacterial pathogen under plant immunity. Proceedings of the National Academy of Sciences 115, E3055–E3064.

O’Connell, R., Herbert, C., Sreenivasaprasad, S., Khatib, M., Esquerré-Tugayé, M.-T., and Dumas, B. (2004). A novel Arabidopsis-Colletotrichum pathosystem for the molecular dissection of plant-fungal interactions. Molecular plant-microbe interactions 17, 272–282.

O’Connell, R.J., Thon, M.R., Hacquard, S., Amyotte, S.G., Kleemann, J., Torres, M.F., Damm, U., Buiate, E.A., Epstein, L., and Alkan, N. (2012). Lifestyle transitions in plant pathogenic Colletotrichum fungi deciphered by genome and transcriptome analyses. Nature genetics 44, 1060–1065.

Ökmen, B., and Doehlemann, G. (2014). Inside plant: biotrophic strategies to modulate host immunity and metabolism. Current opinion in plant biology 20, 19–25.

Osuna-Cruz, C.M., Paytuvi-Gallart, A., Di Donato, A., Sundesha, V., Andolfo, G., Aiese Cigliano, R., Sanseverino, W., and Ercolano, M.R. (2018). PRGdb 3.0: a comprehensive platform for prediction and analysis of plant disease resistance genes. Nucleic acids research 46, D1197–D1201.

Poncini, L., Wyrsch, I., Dénervaud Tendon, V., Vorley, T., Boller, T., Geldner, N., Métraux, J.-P., and Lehmann, S. (2017). In roots of Arabidopsis thaliana, the damage-associated molecular pattern AtPep1 is a stronger elicitor of immune signalling than flg22 or the chitin heptamer. PloS one 12, e0185808.

Rich-Griffin, C., Eichmann, R., Reitz, M.U., Hermann, S., Woolley-Allen, K., Brown, P.E., Wiwatdirekkul, K., Esteban, E., Pasha, A., and Kogel, K.-H. (2020). Regulation of cell type-specific immunity networks in Arabidopsis roots. Plant Cell 32, 2742–2762.

Roussin-Léveillée, C., Lajeunesse, G., St-Amand, M., Veerapen, V.P., Silva-Martins, G., Nomura, K., Brassard, S., Bolaji, A., He, S.Y., and Moffett, P. (2022). Evolutionarily conserved bacterial effectors hijack abscisic acid signaling to induce an aqueous environment in the apoplast. Cell Host & Microbe 30, 489–501. e484.

Ryder, L.S., Cruz-Mireles, N., Molinari, C., Eisermann, I., Eseola, A.B., and Talbot, N.J. (2022). The appressorium at a glance. Journal of Cell Science 135, jcs259857.

Saelens, W., Cannoodt, R., Todorov, H., and Saeys, Y. (2019). A comparison of single-cell trajectory inference methods. Nature biotechnology 37, 547–554.

Sai, N., Bockman, J.P., Chen, H., Watson-Haigh, N., Xu, B., Feng, X., Piechatzek, A., Shen, C., and Gilliham, M. (2022). SAI: Fast and automated quantification of stomatal parameters on microscope images. bioRxiv, 2022.2002. 2007.479482.

Shahan, R., Hsu, C.-W., Nolan, T.M., Cole, B.J., Taylor, I.W., Greenstreet, L., Zhang, S., Afanassiev, A., Vlot, A.H.C., and Schiebinger, G. (2022). A single-cell Arabidopsis root atlas reveals developmental trajectories in wild-type and cell identity mutants. Developmental cell 57, 543–560. e549.

Soneson, C., Love, M.I., and Robinson, M.D. (2015). Differential analyses for RNA-seq: transcript-level estimates improve gene-level inferences. F1000Research 4.

Spanu, P.D., Abbott, J.C., Amselem, J., Burgis, T.A., Soanes, D.M., Stüber, K., Loren van Themaat, E.V., Brown, J.K., Butcher, S.A., and Gurr, S.J. (2010). Genome expansion and gene loss in powdery mildew fungi reveal tradeoffs in extreme parasitism. Science 330, 1543–1546.

Sugio, A., MacLean, A.M., Kingdom, H.N., Grieve, V.M., Manimekalai, R., and Hogenhout, S.A. (2011). Diverse targets of phytoplasma effectors: from plant development to defense against insects. Annual review of phytopathology 49, 175–195.

Takken, F., and Rep, M. (2010). The arms race between tomato and Fusarium oxysporum. Molecular plant pathology 11, 309–314.

Tang, W., Coughlan, S., Crane, E., Beatty, M., and Duvick, J. (2006). The application of laser microdissection to in planta gene expression profiling of the maize anthracnose stalk rot fungus Colletotrichum graminicola. Molecular Plant-Microbe Interactions 19, 1240–1250.

Tian, H., Wu, Z., Chen, S., Ao, K., Huang, W., Yaghmaiean, H., Sun, T., Xu, F., Zhang, Y., and Wang, S. (2021). Activation of TIR signalling boosts pattern-triggered immunity. Nature 598, 500–503.

Trapnell, C., Cacchiarelli, D., Grimsby, J., Pokharel, P., Li, S., Morse, M., Lennon, N.J., Livak, K.J., Mikkelsen, T.S., and Rinn, J.L. (2014). The dynamics and regulators of cell fate decisions are revealed by pseudotemporal ordering of single cells. Nature biotechnology 32, 381–386.

Tsuda, K., and Katagiri, F. (2010). Comparing signaling mechanisms engaged in pattern-triggered and effector-triggered immunity. Current opinion in plant biology 13, 459–465.

Tsuda, K., and Somssich, I.E. (2015). Transcriptional networks in plant immunity. New Phytologist 206, 932–947.

Wang, N., Pierson, E.A., Setubal, J.C., Xu, J., Levy, J.G., Zhang, Y., Li, J., Rangel, L.T., and Martins Jr, J. (2017). The Candidatus Liberibacter–host interface: insights into pathogenesis mechanisms and disease control. Annual review of phytopathology 55, 451–482.

Wang, Y., Bouwmeester, K., Van de Mortel, J.E., Shan, W., and Govers, F. (2013). A novel A rabidopsis–oomycete pathosystem: differential interactions with P hytophthora capsici reveal a role for camalexin, indole glucosinolates and salicylic acid in defence. Plant, cell & environment 36, 1192–1203.

Widemann, E., Bruinsma, K., Walshe-Roussel, B., Rioja, C., Arbona, V., Saha, R.K., Letwin, D., Zhurov, V., Gómez-Cadenas, A., and Bernards, M.A. (2021). Multiple indole glucosinolates and myrosinases defend Arabidopsis against Tetranychus urticae herbivory. Plant Physiology 187, 116–132.

Wilson, R.A., and Talbot, N.J. (2009). Under pressure: investigating the biology of plant infection by Magnaporthe oryzae. Nature Reviews Microbiology 7, 185–195.

Wu, T., Hu, E., Xu, S., Chen, M., Guo, P., Dai, Z., Feng, T., Zhou, L., Tang, W., and Zhan, L. (2021). clusterProfiler 4.0: A universal enrichment tool for interpreting omics data. The Innovation 2, 100141.

Xu, J., Meng, J., Meng, X., Zhao, Y., Liu, J., Sun, T., Liu, Y., Wang, Q., and Zhang, S. (2016). Pathogen-responsive MPK3 and MPK6 reprogram the biosynthesis of indole glucosinolates and their derivatives in Arabidopsis immunity. The Plant Cell 28, 1144-1162.

Xu, Z., Wang, Q., Zhu, X., Wang, G., Qin, Y., Ding, F., Tu, L., Daniell, H., Zhang, X., and Jin, S. (2022). Plant Single Cell Transcriptome Hub (PsctH): an integrated online tool to explore the plant single−cell transcriptome landscape. Plant Biotechnology Journal 20, 10–12.

Yadeta, K.A., and J. Thomma, B.P. (2013). The xylem as battleground for plant hosts and vascular wilt pathogens. Frontiers in plant science 4, 97.

Yang, L., Zhang, Y., Guan, R., Li, S., Xu, X., Zhang, S., and Xu, J. (2020). Co−regulation of indole glucosinolates and camalexin biosynthesis by CPK5/CPK6 and MPK3/MPK6 signaling pathways. Journal of Integrative Plant Biology 62, 1780–1796.

Ye, Q., Zhu, F., Sun, F., Wang, T.-C., Wu, J., Liu, P., Shen, C., Dong, J., and Wang, T. (2022). Differentiation trajectories and biofunctions of symbiotic and un-symbiotic fate cells in root nodules of Medicago truncatula. Molecular Plant 15, 1852–1867.

Yoo, S.-D., Cho, Y.-H., and Sheen, J. (2007). Arabidopsis mesophyll protoplasts: a versatile cell system for transient gene expression analysis. Nature protocols 2, 1565–1572.

Zhang, J., Coaker, G., Zhou, J.-M., and Dong, X. (2020). Plant immune mechanisms: from reductionistic to holistic points of view. Molecular plant 13, 1358–1378.

Zhou, F., Emonet, A., Tendon, V.D., Marhavy, P., Wu, D., Lahaye, T., and Geldner, N. (2020). Co-incidence of damage and microbial patterns controls localized immune responses in roots. Cell 180, 440–453. e418.

Zhu, J., Lolle, S., Tang, A., Guel, B., Kvikto, B., Cole, B., and Coaker, G. (2022). Single-cell profiling of complex plant responses to Pseudomonas syringae infection. bioRxiv.

